# Genomic bases of short-term evolution in the wild revealed by long-term monitoring and population-scale sequencing

**DOI:** 10.1101/2025.10.01.679793

**Authors:** Tristan Cumer, Alexandros Topaloudis, Eleonore Lavanchy, Anne-Lyse Ducrest, Sonia Sarmiento Cabello, Anna Hewett, Paolo Becciu, Céline Simon, Bettina Almasi, Alexandre Roulin, Jérôme Goudet

**Affiliations:** Department of Ecology and Evolution, University of Lausanne, CH 1015, Lausanne, Switzerland; Swiss Institute of Bioinformatics, Lausanne 1011, Switzerland; Swiss Ornithological Institute, Seerose 1, Sempach, CH 6204, Switzerland

## Abstract

Understanding how wild populations adapt to rapid environmental change requires linking phenotypic evolution to its genomic basis over contemporary timescales. This remains challenging because genetic and environmental effects are often intertwined. Here, we leverage a 30-year study of Swiss barn owls (Tyto alba) to explore this process. During this period, owls have evolved darker plumage and increased spottiness, two melanin-based traits associated with fitness. Whole-genome sequencing of 3,102 individuals reveals that these traits are largely controlled by few loci of major effect with partially overlapping architectures. Temporal allele frequency analyses show subtle but consistent shifts at these loci. Simulations indicate that genetic drift alone cannot explain these changes, whereas models incorporating selection do. Our findings demonstrate that selection on a small number of loci can drive rapid phenotypic evolution in the wild. This work underscores the adaptive potential of natural populations and the value of long-term genomic monitoring under accelerating climate change.

## Introduction

Understanding how populations evolve, what mechanisms drive micro-evolutionary changes in phenotypic traits, and how genetic variation translates into variation in fitness are central questions in evolutionary biology. These questions have intrigued evolutionary biology for decades ^1–4^. Addressing these questions is not only essential to evolutionary theory but also critical for understanding how populations can rapidly adapt to changing environments. Achieving such comprehensive understanding of short-term evolution requires detailed knowledge of the links between genotypes, phenotypes, and fitness ^5–7^.

Evolutionary theory predicts that genetically encoded traits evolve toward their optimal values through natural selection, favoring individuals best suited to their environments ^1,3^. In practice, however, identifying adaptive phenotypic change across generations and linking it to the underlying genotypes remains a major challenge. Traditional quantitative genetics approaches can detect heritable changes in life history traits^8^ but often fail to pinpoint specific causal loci. Conversely, population genetics approaches can identify genomic regions associated with phenotypic traits^9^, though their application to wild, non-model species has been limited by cost and logistical constraints. Recent advances in whole-genome sequencing and downstream analyses ^10^ now enable the study of large cohorts in wild populations ^11,12^. By combining quantitative genetic methods with high-quality genomic data, we can explore the genomic basis of phenotypic evolution in the wild with unprecedented resolution.

Among phenotypic traits, coloration offers a powerful system for studying the genotype-phenotype fitness relationship, as well as their joint evolution ^13^. Extensive research on the genetic basis of animal coloration spans humans ^14–16^, model organisms ^17–19^ and non-model species ^20–23^, yielding deep insight into both the molecular and evolutionary mechanisms of color variation ^24,25^. However, most studies focus on discrete color polymorphisms ^20–23,25^ while continuously varying color traits, which are common in natural populations, remain underexplored ^24,25^. This is largely due to the challenges of disentangling environmental and genetic influences on continuous traits, which often result from numerous variants, ranging from large to small effect sizes, frequently located in regulatory regions ^24–26^ (with a few exceptions ^27–29^). Yet, these traits are evolutionary relevant because coloration often directly influences fitness by affecting courtship, predator-prey interactions, thermoregulation, immunity, and other ecological interactions ^30^. Consequently, an individual’s color is commonly under multiple selective pressures, often related to both interspecific and intraspecific interaction ^13,31^.

Color polymorphism in the barn owl (*Tyto alba*) offers a particularly suitable system to explore the genomic basis of fitness-related trait. In Central Europe, barn owls exhibit continuous variation in two melanin-based traits on the ventral body: plumage coloration (ranging from white to rufous) and spottiness (number and size of black feather spots) (Figure 1A). Both traits are related to fitness: plumage coloration is linked to fecundity ^32,33^ and hunting success ^34,35^. Spottiness, in turn, is related to testosterone levels ^36^, stress response ^37,38^ and immunity ^39,40^. Both traits are heritable, and a genetic correlation between them suggests shared genomic underpinning ^41^. Females tend to be more melanic (redder and spottier) than males ^42^, and spottiness shows higher heritability in males ^41^ suggesting potential sex-linked genetic effects ^43^.

**Figure 1:**
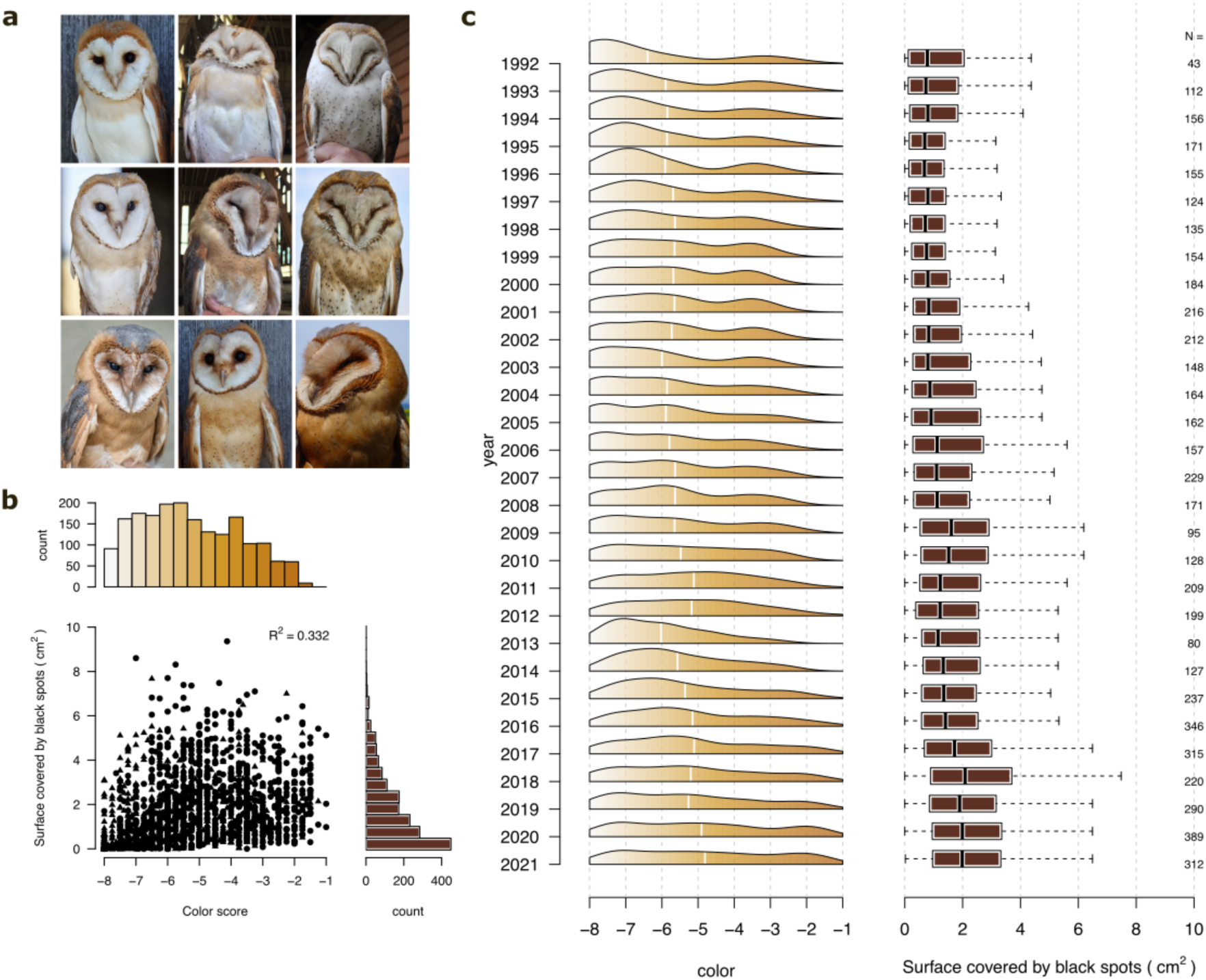
Variation in plumage coloration and spottiness of Swiss barn owls over the last 30 years. (a) Photographs illustrating the range of ventral plumage coloration (rows) and spottiness (columns) observed in Swiss barn owls. Coloration varies from white (-8) to dark reddish (-1), while spottiness reflects variation in the density and size of black spots. (b) Relationship between coloration and spottiness (in cm^2^). The upper panel shows histograms of the distribution of coloration, while the right panel shows the distribution of spottiness. (c) Temporal trends in coloration and spottiness in alive adults between 1992 and 2021. Left: density plots of coloration for each year, with the white line indicating the annual mean. Right: boxplots of spottiness for all individuals alive in each year; sample sizes are reported on the right.

A recent continental-scale GWAS identified two loci associated with plumage coloration: a major SNP in the *MC1R* gene on chromosome 11 (autosome), and a second locus on the Z sex chromosome ^29^. While these loci explain much of the variation in coloration, ∼20% remains unexplained. Moreover, the genetic basis of spottiness is still unexplored. This gap highlights the importance of investigating whether these two traits share common genetic determinants, if sex-specific effects contribute to their architecture, and how these factors shape their evolutionary trajectories.

Taking advantage of over 30 years of phenotypic and life-history monitoring in a wild barn owl population in western Switzerland, we investigated the microevolutionary dynamics of coloration and spottiness. Over this period, we observed a clear phenotypic trend toward redder and more spotted individuals. To uncover the genetic drivers of this trend, we sequenced 3102 individuals, improving resolution of genetic relatedness and refining heritability estimates. We identified three causal loci associated with these traits, including a shared signal on the Z chromosome. Gene expression analyses confirmed the link between Z-linked genotypes and phenotypes, suggesting a mechanistic basis for the observed sexual dimorphism. We further modelled the sex-specific genetic architecture of the traits, revealing differing allelic effects between males and females. Finally, forward simulations showed that selection, rather than genetic drift alone, can explain the observed allele frequency shifts.

## Results and discussion

### Phenotypic changes in the two melanic traits

In the Swiss barn owl population, plumage color and spottiness show substantial variation (Fig. 1a, 1b). Color scores range from-8 (white) to-1 (red), with a mean of-5.21 (SD=1.78; Fig. S1), while spottiness, measured as the area covered by black feather spots, average 1.71cm^2^ (SD=1.54; Fig. S1; n= 8278). The two traits are moderately correlated (Pearson’s r 0.33; Fig. 1b, Fig. S2-S3), with a significantly stronger correlation in males (r=0.37) than in females (r= 0.18; Fig. S2).

Phenotypic data collected over the 30-year study period reveal a clear evolutionary trend toward redder and more heavily spotted individuals (Fig. 1c). Linear mixed-effects models confirm a significant temporal increase in both traits: color scores increased by 0.04 units per year, and spottiness increased by 0.05 cm^2^ per year. Over 30 years, these changes represent shifts of 0.66 and 0.91 standard deviations from the respective means (Tables S1-S2 for color; Tables S3-S4 for spottiness). This trend was consistent across both males and females showing increased melanism over time (Fig. S4-S7, Tables S5-S8 for sex-specific models), despite the species’ weak sexual dimorphism (Fig. S6). Moreover, the trend held when separating individuals born within the study area (residents) from those born outside it (immigrants), indicating that it is not driven by selective migration (Figs. S8-S9). Together, these results show that both coloration and spottiness have undergone directional change in this population, with barn owls becoming progressively redder and more spotted over the past three decades.

### Population-scale whole-genome sequencing of the Swiss barn owl population

To investigate the genetic architecture underlying variation in coloration and spottiness, we assembled a whole-genome dataset consisting of 3,102 barn owls from the Swiss population, sampled over a 31-year period. We leveraged a previously published high-coverage dataset of 502 owls from across the Western Palearctic, including 346 from our focal Swiss population, to construct a reference panel representing continental genomic diversity (10,451,268 phased SNPs; Fig. S10–S13; ^44^). Using this panel, we imputed genotypes for an additional 2,778 Swiss individuals sequenced at low coverage (mean 1.95X; Fig. S14), achieving high imputation accuracy (mean r^2^=0.968). The final dataset comprised 3,280 individuals of which 3,102 were phenotyped for coloration and spottiness, providing an unprecedented resource for dissecting the genomic basis and evolutionary dynamics of these traits.

### High heritability and large-effect loci shared between the two traits

Using whole genome data from 3102 owls, we first confirmed the high heritability of both plumage coloration and spottiness. Genetic relatedness was estimated using a genomic relationship matrix (GRM) derived from SNP data, and heritability was estimated using an animal model. Coloration showed a heritability of h^2^= 0.82 ± 0.02 (Table S9), consistent with previous estimates based on pedigree data (h^2^= 0.84 ± 0.02) ^41^. Spottiness exhibited similarly high heritability (h^2^=0.80 ± 0.02; Table S10), exceeding prior estimates for its two components-number of spots (h^2^= 0.57 ± 0.03) and spot diameter (h^2^= 0.67 ± 0.03) ^41^ - likely due to differences in how the trait was measured. Unlike previous analyses treating the components separately, our approach models total spottiness as an integrated phenotype. Together, these results demonstrate a strong genetic basis for both traits.

We next estimated the genetic correlation between coloration and spottiness to assess the degree of shared genetic architecture. The overall genetic correlation was moderate (r = 0.24 ± 0.08), indicating partial but not complete genetic overlap. However, this correlation was substantially higher in males (r = 0.55 ± 0.11) than in females (r = 0.11 ± 0.16), suggesting sex-specific differences in genetic control. These results, consistent with previous findings in barn owls ^41^, support the involvement of sex-linked genetic factors, particularly variation on the Z chromosomes, in shaping these traits.

### Genome-wide association studies reveal shared and distinct large-effect loci

To identify genomic regions associated with coloration and spottiness, we performed independent genome-wide association studies (GWAS) for each trait (Fig. 2, S15-S20; Table S11-S20). For coloration, we identified two significant loci (after Bonferroni threshold:-log_10_(p> 8.11; Fig. 2a, 2c and 2d). The top signal was on chromosome 11 (Super-scaffold 26) at position 22,522,039, a known nonsynonymous variant in the *MC1R* gene ^45,46^. This variant is referred to as MC1R_white_ (G) and MC1R_red_ (A) based on associated phenotypes. A second peak was located on the sex chromosome (chromosome Z - Super-Scaffold 42) in a region (between 29’200’000 and 30’200’000 bp) encompassing 16 genes, with the most associated SNP on position 29’826’792 (Fig. 2a and 2d, Table S11). This variant is referred to as Z_white_ (G) and Z_red_ (C), based on associated phenotypes. Both loci confirm associations previously reported at the continental scale ^46^. To account for spatial heterogeneity in coloration across the ventral surface, we ran additional GWAS separately for breast and belly coloration (Fig. S16, S17, Table S12, S13). The breast-specific GWAS revealed an additional locus on chromosome 4 (Super-Scaffold 6) at 27,439,651 bp, located within an intron of the *CORIN* gene (-log_10_ (p)=9.43), not detected in the full-body analysis.

**Figure 2:**
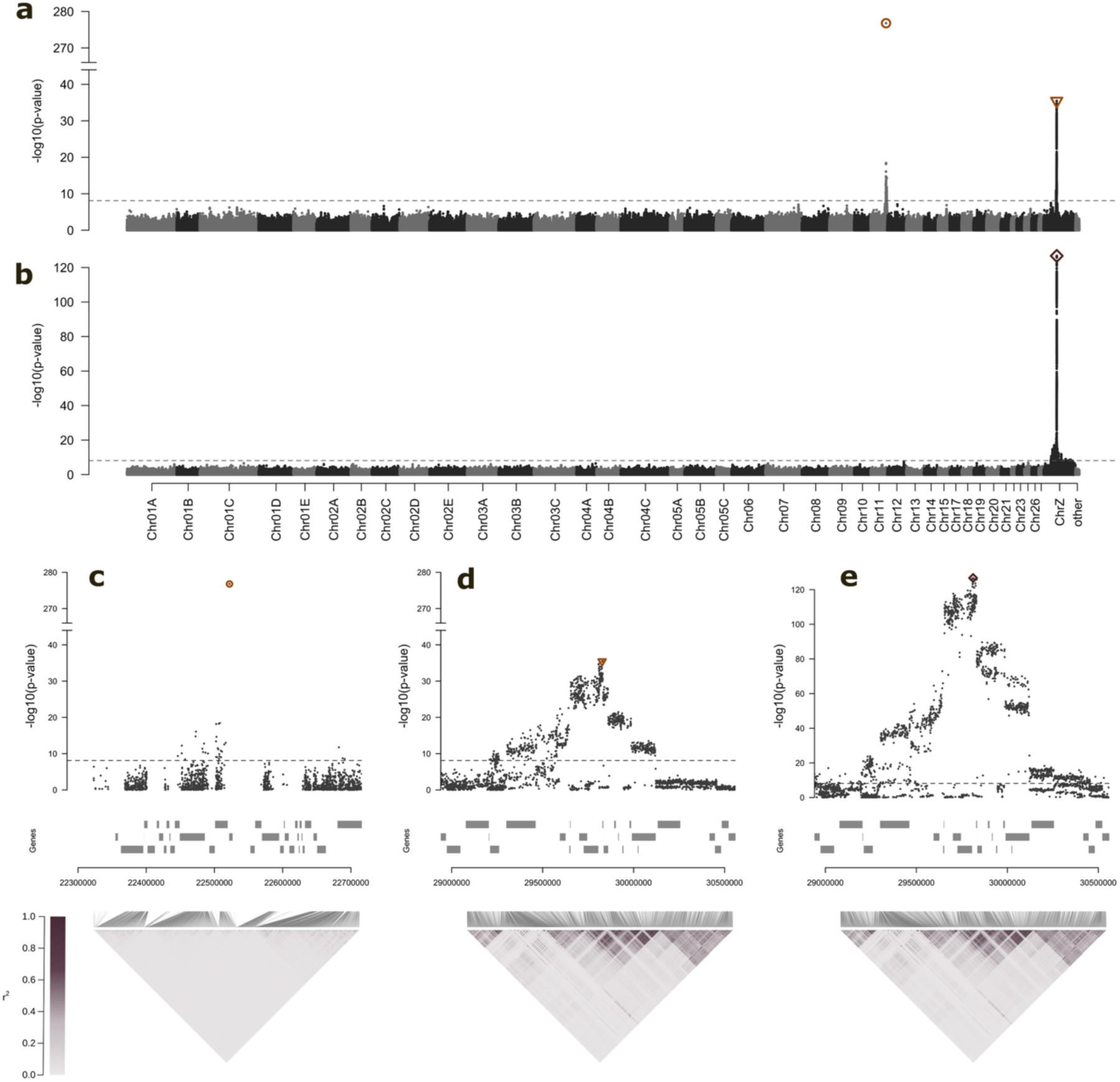
Genome-wide association study (GWAS) of plumage traits in Swiss barn owls reveals an oligogenic architecture for the two traits. Association scores (–log₁₀ p-values) for each SNP across the genome with (a) plumage coloration and (b) plumage spottiness. Alternating colors distinguish successive chromosomes. The dotted line indicates the Bonferroni-corrected significance threshold (-log^10^(p-value) = 8.11). circle, triangle and diamond point to the most significant SNP in each region presented in (c–e). (c–e) Zoomed views of regions showing strong associations with coloration (c and d) or spottiness (e). Upper panels display association scores for SNPs within each region, with dotted lines marking the Bonferroni threshold. Rectangles below indicate annotated genes. Lower panels show mean linkage disequilibrium (r²) between SNPs in sliding windows of 10 SNPs. Lines connect each LD block to its genomic position on the x-axis.

For spottiness, the GWAS revealed a single significant region on the Z chromosome, overlapping with the region associated with coloration (Fig. 2b, 2e). The top SNP was located at position 29’811’663 (-log10(p< 8.11; Fig 2b, 2e). The associated alleles are referred to as Z_immaculate_ (G) and Z_spotted_ (A), based on phenotype. To further dissect spottiness, we conducted a GWASs on its two components, mean spot diameter and total number of spots. The GWAS on spot diameter showed a pattern similar to the total surface area, with significant associations primarily on the Z chromosome (Fig. S19, S20, Table S15, S16). In contrast, the GWAS for number of spots identified multiple significant regions, including the MC1R variant on chromosome 11.

The GWAS results indicate an oligogenic architecture for both traits, with a few large-effect loci driving much of the variation. Notably, both traits share a common genetic architecture, with overlapping signals at the Z chromosome and the *MC1R* gene: (i) There was almost complete linkage disequilibrium (R²=0.934) between the top SNPs for Z coloration and Z spottiness, indicating near-identical genotype-phenotype associations; (ii) The MC1R locus classically known for its role in pigmentation across vertebrates, also contributes to variation in both color and spot number. By precisely localizing the shared region on the Z chromosome and confirming its role in both traits, our work results refine previous estimates of the genetic correlation between coloration and spottiness ^41^. Interestingly, this Z-linked region has also been associated with color variation in North American rosy finches ^47^, suggesting a potentially conserved pigmentation pathway across species.

### Differential gene expression links genotype to phenotype

To investigate the molecular mechanisms linking genotype and phenotype, we measured the expression of genes within the GWAS-identified regions. Specifically, we quantified expression of 13 candidate genes-*MC1R* and 12 genes located in the Z-linked region associated with both traits (Fig. S21, S22, Table S17), in developing breast feathers from 45 nestlings (28 males and 17 females). Sampling was timed to capture the onset of feather development, when black apical spots and pheomelanin-based coloration first become visible.

The nonsynonymous MC1R (V126I) has previously been associated with both coloration and spot number, as well as with variation in *MC1R* gene expression ^48^. We included MC1R in our expression panel, but due to a sampling strategy designed to maximize Z-linked genotype variation, the resulting MC1R genotype distribution was highly unbalanced (38 homozygous MC1R_white_, 6 heterozygous and 1 homozygous MC1R_red_), limited statistical power to assess its regulatory effect. We thus focused on the Z-linked region, where we quantified the relative expression levels of all 12 genes within the top GWAS peak (Fig. 3a). First, we confirmed that genotypes at the top Z-linked SNP were significantly associated with both coloration and spottiness in these individuals (Fig. 3b, 3c). We then used a linear mixed model to test whether gene expression levels varied with genotype at the Z locus. As expected, expression levels differed significantly between sexes for all 12 genes, typically higher in males, consistent with incomplete dosage compensation in birds ^49–51^. To account for this, we ran sex-specific models to test for genotype-expression associations (Table S18). Of the 12 genes tested, two showed significant genotype-dependent expression: *PDE8B* varied significantly with genotype in males, with lower expression in heterozygotes (Fig. 3d, Table S18-19), *WDR41* with genotype in females (Fig. 3e, Table S21-22). To assess whether these expression differences translated into phenotypic effects, we tested for associations between expression and trait values. Only one significant link emerged with PDE8 expression in males associated with spottiness, but not coloration (Fig. 3f, 3g, S22, Table S23)

**Figure 3:**
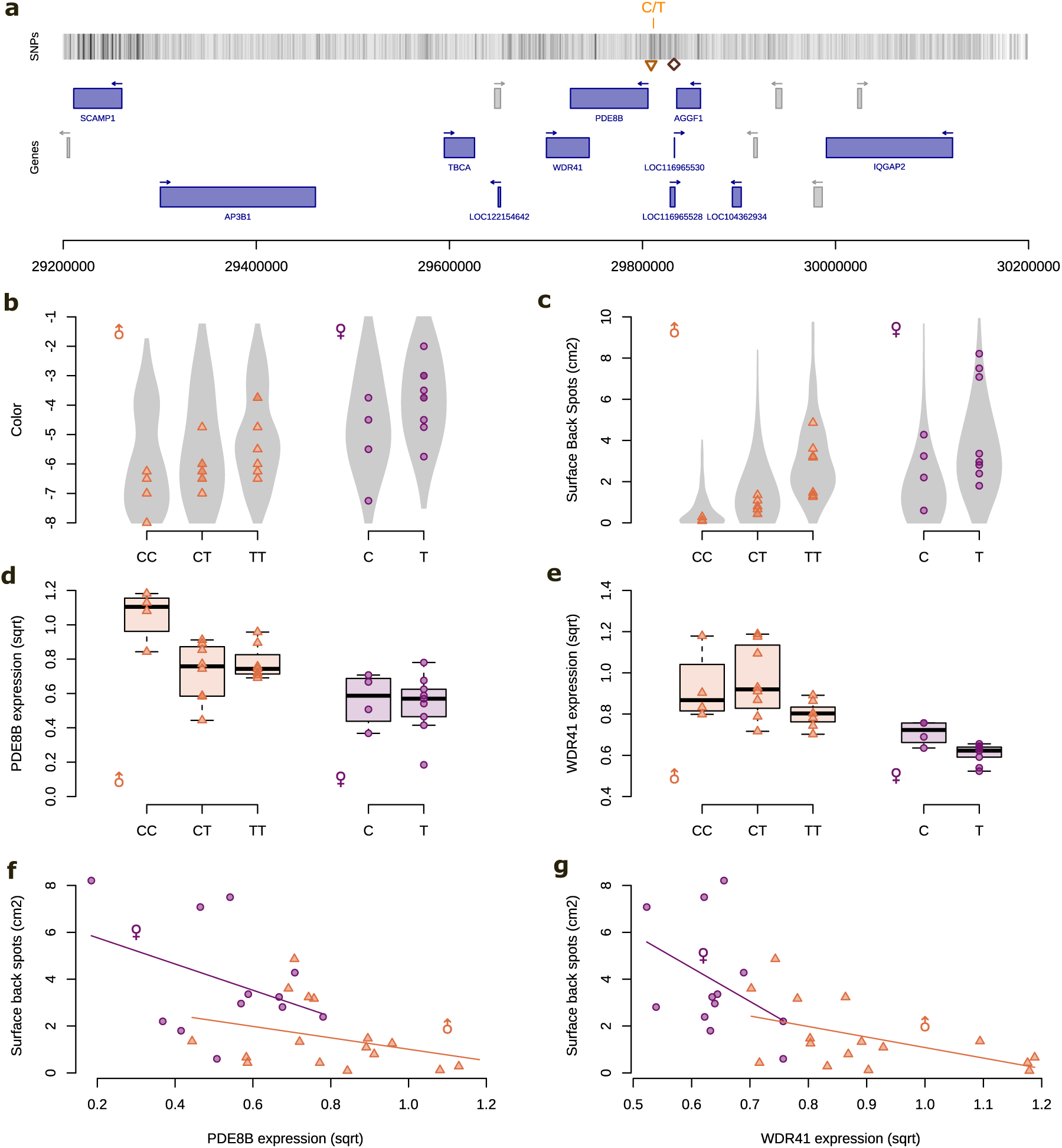
Candidate genomic region and genotype–phenotype associations for plumage traits in Swiss barn owls. (a) Genomic region on chromosome Z (Super Scaffold 42) containing annotated genes, including PDE8B and WDR41. The focal SNP (C/T) and the most significant SNP for color and spottiness are indicated. (b,c) Violin plots of plumage color and spottiness (black spot area, cm²) across genotypes (CC, CT, TT in males; C, T in females) for 3,102 sequenced individuals. Triangles and circles denote the 45 individuals with quantified gene expression. (d,e) Boxplots of sqrt-transformed expression of PDE8B and WDR41 across genotypes. (f,g) Scatter plots showing correlations between sqrt-transformed expression and spottiness for PDE8B and WDR41; lines depict sex-specific linear models.

Our findings suggest a potential link between genotype, gene expression, and the spottiness phenotype in barn owls. However, this association remains preliminary and warrants further investigation. One limitation of our study is the small sample size, which prevented us from accounting for variation in the *MC1R* genotype, potentially obscuring true genetic associations. Despite this constraint, we identified *PDE8B* as a promising candidate gene. *PDE8B* encodes a cAMP-specific phosphodiesterase ^52^, and although it has not previously been implicated in pigmentation, other phosphodiesterases, such as PDE5, are known to influence melanin synthesis via the cAMP/PKA/CREB/MITF signaling pathway ^53,53,54^. Further research is needed to clarify the molecular role of the Z-linked region. Given its location on the sex chromosome and the pronounced sexual dimorphism in barn owls, we also considered whether sex-specific genetic architecture might contribute to variation in these traits.

### Sexual dimorphism and sex specific genetic architecture of the two traits

The barn owl is a sexually dimorphic species, with males typically exhibiting whiter and fewer ventral spots than females ^55^. To investigate how the genomic variants identified in our GWAS contribute to phenotypic variation in each sex, and to quantify the relative effect of each locus, we fitted Bayesian generalized linear models separately for each trait. Genotype at each locus was fitted as fixed effects. Given the pronounced sexual dimorphism and the fact that one of the loci most strongly associated with both traits is located on the Z chromosome, we constructed sex-specific models. In total, we ran four models, one for each trait (coloration and spottiness) and for each sex (Figure 4). To control for ontogenic changes in phenotype, we included the developmental stage of each individual (fledgling or adult) as covariate in all models ^56^.

**Figure 4:**
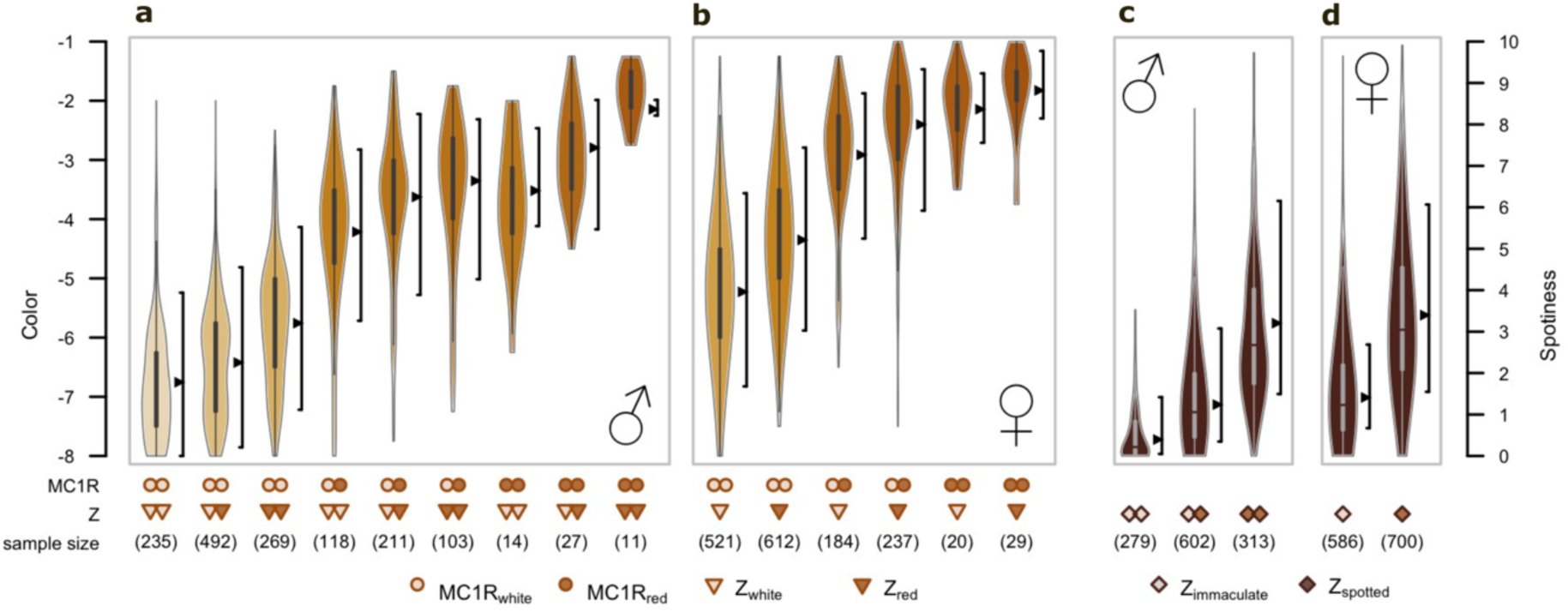
Sex-specific genetic architecture of plumage coloration and spottiness in Swiss barn owls. (a-b) Violin plots of observed plumage coloration across genotype combinations for males (a) and females (b). Violin colors indicate mean phenotype per genotype. Intervals to the right of each violin show predicted lower and upper whiskers from the genetic architecture model; black triangles indicate median predicted values. The x-axis shows genotype dosage for each locus; sample sizes are in parentheses. (c) Same as (d), for plumage spottiness.

The two sex-specific models for plumage coloration confirm the contribution of three loci-*MC1R*, the Z-linked region, and *CORIN*)-in both males and females. The models explained 80.7% (± 1.6) of the total variance in coloration in males and 75.3% (± 2.1) in females (Figure 4a, b). These values closely match the previously estimated heritability of the trait (0.82 ± 0.02; see *High heritability and large-effect loci shared between the two traits* section), supporting the interpretation of plumage color as an oligogenic trait with most of the variance accounted for by few loci. Genotypes at all three loci had significant effect sizes (Table S24-S27). Consistent with earlier findings, *MC1R* emerged as the major contributor to color variation ^46,57^), followed by the Z-linked locus ^46^. The significant effect of *CORIN*, however, represents a novel result. Its detection here highlights how increasing sample size can reveal loci with smaller effects in oligogenic traits. While all three loci influenced coloration in both sexes, the magnitude and nature of their effects differed. Notably, a significant epistatic interaction between *MC1R* and the Z-linked genotype was detected only in females (Tables S27). In females, the accumulation of “red” alleles at *MC1R* and the Z-locus did not result in a linear increase in redness. Instead, coloration reached a plateau (Fig. 4b), consistent with a non-additive (epistatic) relationship between these two major pigmentation genes. This epistatic interaction has been previously hypothesized ^57^ and is now proven using genomic data.

To model the contribution of the two loci, *MC1R* and the Z-linked region, to spottiness in each sex we used a generalized linear model with a hurdle-gamma distribution. This modelling approach separates (i) the effect of each locus on the presence or absence of spots (the hurdle part, estimating the probability that spottiness equals zero) and (ii) the contribution on the total surface area covered by spots in individuals using the gamma part of the model ^58^. In males, both loci significantly contributed to the presence / absence of spots. However, the Z-linked locus consistently had a larger effect than *MC1R* on both the likelihood of being spotted and the extent of spottiness (Figure 4c, see table S28-S31). In females, while the Z-linked locus had a significant impact on both the presence of spots and the total spotted surface, the effect of *MC1R* was not statistically significant (Figure 4d, Table S31). These results suggest a slightly different architecture for spottiness between the sexes. The models explained 58.3% (±2.5) and 53.9% (±3.1) of the variance in spottiness in males and females, respectively. While this represents a substantial proportion of the trait variance, it falls short of the higher heritability previously estimated for spottiness (0.80 ± 0.02; see *High heritability and large-effect loci shared between the two traits* section). This discrepancy indicates that additional genetic or environmental factors likely contribute to the trait.

Taken together, our results reveal distinct genetic architectures for both coloration and spottiness between the sexes. The epistatic interaction between *MC1R* and the Z-linked locus in determining coloration in females, alongside the male-specific contribution of *MC1R* to spottiness, suggests that trait expression is influenced not only by individual loci but also by their interactions, interactions that may differ between sexes with mechanisms such as incomplete dosage compensation in birds ^49–51^ and sex-biased gene expression ^59,60^, both of which can reduce antagonistic sexual conflict ^61^. These findings underscore the importance of modelling sex-specific genetic effects and inter-locus interactions when investigating the evolution of complex traits and the maintenance of sexual dimorphism in natural populations.

Given the large effects of these loci on phenotype and the observed changes in trait distribution over time, we next leveraged the temporal dimension of our dataset to assess whether shifts in allele frequencies could account for the observed phenotypic trends.

### Characterizing the evolutionary processes driving allelic and phenotypic changes

To investigate the evolutionary forces underlying allele frequency changes at the *MC1R* and Z-linked loci, we leveraged the temporal resolution of our 30-year dataset to track allele dynamics over time (Fig. 5). Allele frequencies were estimated annually using genotypes from adult owls alive in each year (Fig. 5b, 5e). Over the three-decade period, the frequency of the *MC1R_red_* allele increased significantly, while the Z_spotted_ allele showed a significant decline (Fig. 5b, 5e, Table S32-S33). Given the strong associations of these loci with plumage coloration and spottiness, respectively, this temporal trend may reflect directional selection acting on these traits. However, change in allele frequencies over time could also be influenced by gene flow, demographic changes, or genetic drift. To disentangle these processes, further analysis is needed to assess the role of selection relative to neutral evolutionary forces.

**Figure 5:**
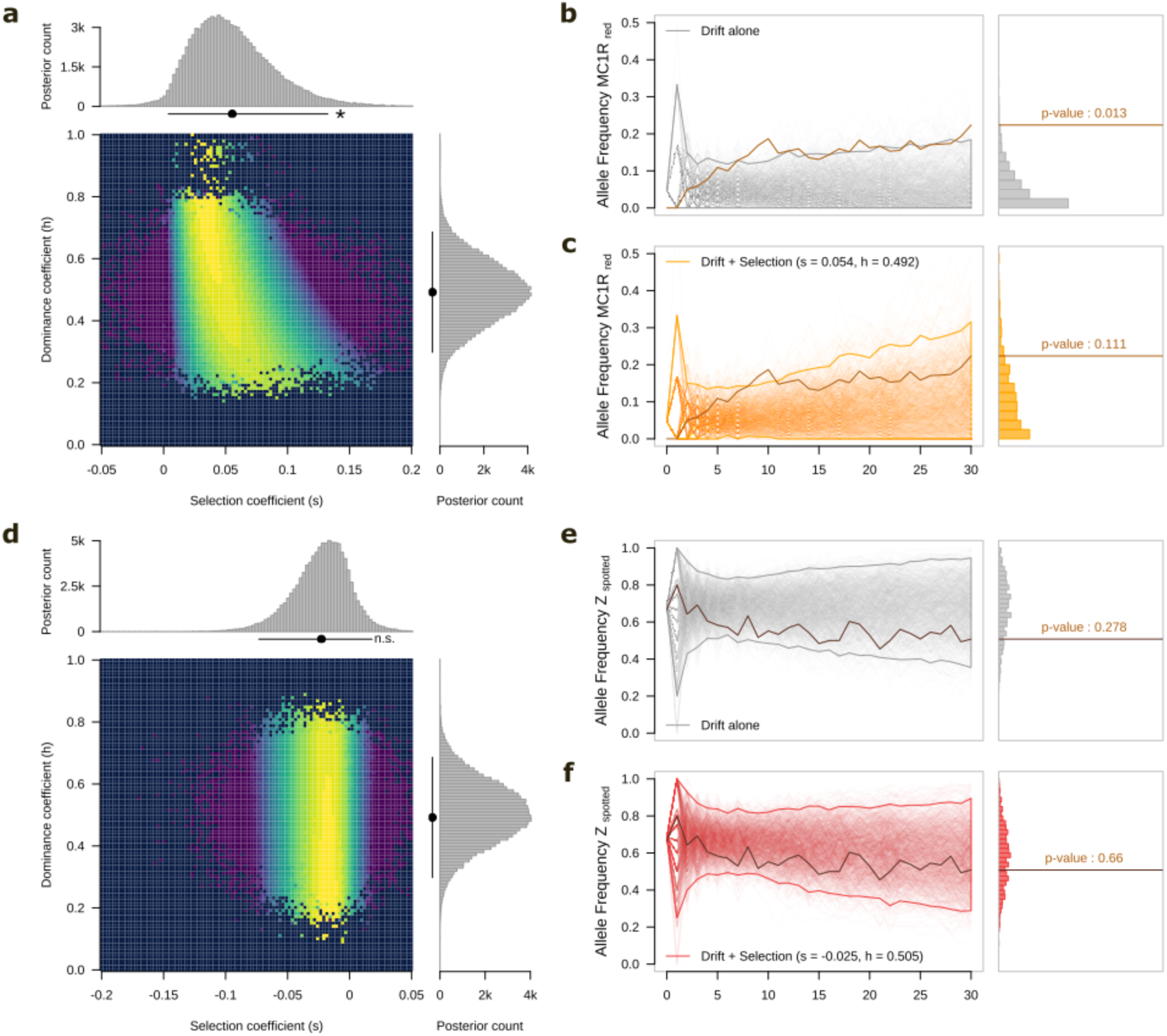
Likelihood-based inference of selection parameters and simulation-based validation for MC1R (top panels) and Z_spot_ (bottom panels). (a, d) Heatmaps show the log-likelihood surface across a grid of selection coefficients (s) and dominance coefficients (h), with warmer colors indicating higher likelihood of the observed allele frequency trajectory for a given parameter combination. Overlaid bar plots represent the posterior distributions of s and h, summarizing parameter uncertainty; points below the histograms mark posterior means, and bars indicate 95% confidence intervals. Significance was assessed by whether the confidence interval overlaps zero. (b, c, e, f) Results from forward simulations of allele frequency change under two evolutionary scenarios: genetic drift alone (top) and drift combined with selection (bottom). Histograms show the distribution of final allele frequencies across replicates, and p-values indicate the probability of observing the empirical data under each scenario.

As a first step in disentangling the processes driving allele frequency shifts, we tested whether migration could explain the observed changes. Individuals ringed as fledglings were classified as residents, while those first captured as adults, likely born outside the study area, were considered immigrants. Across years, allele frequencies did not differ significantly between these two groups (Fig. S23) suggesting that migration is unlikely to be the primary driver of the observed allele frequency changes. Instead, the trends likely reflect broader-scale evolutionary processes acting across the wider population. This interpretation aligns with previous findings of high connectivity and low genetic differentiation among barn owl population across continental Europe ^62^.

To assess whether the observed allele frequency changes could be attributed to selection rather than genetic drift, we considered the relationship between allele effects on fitness and population size. For natural selection to act efficiently, the variant must influence fitness, and the effective population size (N_e_) must be large enough for selection to overcome stochastic fluctuations due to drift. In small populations, drift can dominate, causing allele frequencies to change regardless of selective advantage. Census estimates for the Swiss barn owl population remained relatively stable over the study period, ranging from 200 to 1,000 breeding pairs ^63^. To estimate N_e_, we used whole-genome data to calculate nucleotide diversity (π), applying the equation N_e_ = π/(4*μ), where μ is the mutation rate ^64^. Across different subsampling, N_e_ estimates were highly consistent, averaging 226,051 individuals (range: 223,937 in 2016 to 228,716 in 1992; Fig. S24), roughly 100 times larger than the census population. This inflation likely reflects a large historical effective size and high contemporary connectivity among European barn owl populations (low genome-wide differentiation across the Western Palearctic (overall F_ST_=0.047)) ^62^ and rare occurrence of inbreeding ^65^.

To test whether genetic drift alone could explain the observed allele frequency changes, we simulated allele frequency trajectories under a neutral model using the census population size as input (see Methods). For the *MC1R* locus, the observed frequency trajectory of the *MC1R_red_* allele deviated significantly from the range expected under neutrality. Specifically, the increase in MC1R_red_ frequency exceeded that predicted by drift alone (Fig 5b), suggesting a likely role for positive selection. Although observed trajectory exhibited some year-to-year variability, potentially due to limited sample sizes in certain years, family structure within the dataset, or temporal fluctuations in selection intensity, the overall mismatch between simulated and observed patterns indicates that drift alone cannot account for the changes at this locus. In contrast, the frequency dynamics of the Z spotted allele fell within the distribution of trajectories expected under neutrality drift (Fig. 5e), suggesting that the decline in this allele could plausibly be explained by stochastic processes.

To further investigate whether the observed allele frequency changes at the two loci reflect directional selection, we applied a probabilistic modeling framework that infers evolutionary parameters from temporal frequency shifts (see *Methods* for details).

The MC1R locus exhibited a stronger signal of selection. Although only 25% of iterations in the combined model (which includes both selection and drift) supported selection, the selection-inclusive models consistently yielded higher log-likelihoods than the drift-only model (Fig. S25-S27). Furthermore, both the combined and selection-only models produced selection coefficient estimates whose 95% credible intervals excluded zero (Fig. 5a), indicating a statistically significant deviation from neutrality. These results provide compelling evidence that positive selection contributed to the observed increase in *MC1R_red_* allele frequency. The estimated selection coefficient of s=0.054, with a dominance coefficient of h=0.492, suggesting an additive to slightly dominant effect of the *MC1R_red_* allele.

To validate our selection inference for *MC1R*, we incorporated the estimated genotype-specific selection coefficients into forward-time simulations. Under a model combining genetic drift and directional selection, the observed allele frequency trajectory for the *MC1R_red_* allele fell within the simulated range (Fig. 5), further supporting the of selection in shaping its increase over time. In contrast, the analysis of allele frequency changes at the Z-linked locus revealed limited evidence for selection. In the model allowing both drift and selection, 96.5% of iterations favored the drift-only scenario. The log-likelihood difference between the selection and drift models modest (Fig. S28-S30), and the 95% credible interval for the selection coefficient in the selection-only model included zero (Fig. 5d), indicating no statistically significant deviation from neutrality. Moreover, incorporating the estimated selection coefficient into simulations only marginally improved the fit to the observed frequency trajectory (Fig. 5f). Taken together, these results suggest that the allele frequency dynamics at the *MC1R* locus are best explained by a combination of drift and positive selection, whereas those at the Z locus are consistent with neutral evolution.

## Conclusion

Three decades of phenotypic monitoring combined with whole-genome sequencing of over 3,000 individuals provide an unprecedented window into the genomic basis of rapid trait evolution in a wild vertebrate population. We identify an oligogenic architecture underlying two melanin-based traits—coloration and spottiness—both of which have undergone significant shifts over time. These traits share a partially overlapping genetic basis but exhibit pronounced sex-specific differences in genetic architecture and gene interactions. Allele frequency changes at major loci are consistent with selection, although the specific fitness consequences and potential trade-offs with other traits remain to be explored. Crucially, this study illustrates the power of integrating genomic, phenotypic, and long-term ecological data. Such an approach not only deepens our understanding of how wild populations respond to environmental change but also offers a framework for predicting the pace and mechanisms of evolutionary change in natural systems.

## Material and Method

### Study system and phenotyping

#### Study system

A wild population of barn owls (*Tyto alba*) in western Switzerland (46°49ʹ N, 06°56ʹ E) (*Tyto alba*) has been continuously monitored since 1986, spanning over 30 years. These owls typically reach sexual maturity within their first year of life ^66^, and primarily breed in nest boxes installed on farms. Throughout each breeding season, nest boxes were inspected at four-week intervals to assess occupancy. When an active nest was detected, females were captured during incubation, while males were typically caught later during the provisioning of nestlings. During these visits, all nestlings and any previously unmarked adults were fitted with uniquely numbered metal rings for individual identification. Blood samples were collected from the brachial vein of each bird and initially stored in liquid nitrogen before being transferred to-80°C freezers for long-term preservation. Nestling sex was determined using sex-specific molecular markers targeting the SPINDLIN-gene ^67^. Breeding females were identified by the presence of a brood patch. The year of birth was known for individuals ringed as nestlings. For adults ringed for the first time, age and year of birth were estimated based on molt patterns of wing flight feathers.

#### Phenotype measurements

At each capture event, two melanin-based plumage traits, namely coloration and spottiness, were assessed on both the breast and the belly of each individual. The degree of reddish pheomelanic coloration was scored using an eight-color chip scale ranging from −8 (white) to −1 (dark reddish). Spottiness was quantified by counting the number (N) and measuring the mean diameter (D ± 0.1 mm) of eumelanic black spots within a 60 × 40 mm (2400 mm^2^) frame. For each body part (breast and belly), we calculated the total surface area covered by black spots (referred to as spottiness, S) using the following formula (expressed in cm^2^): S = 100 × (N × (𝜋 × (D/2) ^2^)). The spottiness values of the breast and belly were then summed to obtain a single value representing total the ventral spottiness. To retain a single phenotype value per individual, we averaged trait values across all available adult captures. If no data were available (e.g., due to mortality or dispersal), the fledging measurement was retained. The life stage at which individual was phenotype (fledging or adult) was recorded for downstream analyses. Individuals first ringed as adults were classified as “immigrants”, whereas those ringed as fledglings were classified as “residents”. To examine the relationship between the two traits, we estimated Pearson’s correlation coefficient (r) using the *cor()* function R (v4.2.2) ^68^ across all individuals, and separately for each sex. Differences in trait values between sexes and between life stages (juvenile vs. adult) were tested using the *wilcox.test()* function in R.

### Evolution of two melanin-based traits

#### Modeling the temporal evolution of coloration and spottiness

To assess whether the melanin-based traits of coloration and spottiness have changed over time, we modeled the evolution of each trait across years using linear mixed-effects model (LMM). Models were fitted via restricted maximum likelihood (REML) using the *lmer(*) function from the lme4 package (v1.1-37) ^69^ in R (v4.2.2). For each trait, we constructed a series of increasingly complex models by sequentially adding explanatory variables. Model comparisons were based on Akaike’s Information Criterion (AIC; Table S1, S3). The best supported model for both traits included the following fixed effects: year of birth (continuous) to test for long-term evolutionary trends; sex (categorical) to account for sexual dimorphism in plumage traits ^70,71^, Fig. S6); phenotypic stage (categorial: fledging or adult) to control for age-related changes in melanin expression ^56,57^, Fig. S7). To account for variation related to observation timing, we included year of first capture as a random effect in all models. We also tested the effect of origin (resident versus immigrant) by including a categorical fixed effect indicating whether an individual was born within the study population. However, adding this variable did not improve model performance and was therefore excluded from the final model.

### Whole genome sequencing and Genotyping

#### Reference panel: Data collection, phasing, and quality control

The reference panel used in this study was previously assembled ^44^, and includes samples from previous whole-genome sequencing efforts ^44,46,62,72–74^. In total, whole genome sequencing data from 502 samples were processed through a standardized variant discovery pipeline, briefly summarized below (see ^44^ for further details). Raw sequencing reads were processed with Trimmomatic v0.39 ^75^: sequencing adapters were removed, and reads shorter than 70bp were discarded. Cleaned reads were aligned to the *Tyto alba* reference genome (NCBI RefSeq assembly: GCF_018691265.1) ^73^ using BWA-MEM v0.7.17 ^76^.

Variant discovery followed the GATK v4.2.6 best practices pipeline ^77^. Base Quality Score Recalibration (BQSR) was performed using a previously validated truth set ^62^. SNPs were initially called per individual using HaplotypeCaller, then merged via joint genotyping with *GenotypeGVCFs*. Variants were filtered to retain only high-quality bi-allelic SNPs that passed the following GATK hard filters: QD<2.0, QUAL<30, SOR>3.0, FS>60.0, MQ<40.0, MQRankSum<-12.5, and ReadPosRankSum<-8.0). In addition, regions with low mappability were excluded ^78^. Depth-based filtering was performed using BCFTools (v.1.15.1) ^79^: any genotype with read depth < 5 or > (mean depth + 3 SD) was set to missing. SNPs with a minor allele count (MAC) < 5 were also removed. Following all filtering steps, the final reference panel comprised 28’197’066 high-confidence SNPs. To facilitate downstream analyses, SNPs positions originally mapped to Super-scaffolds were converted to chromosomal coordinates based on the linkage map of the barn owl ^44^. Of the retained SNPs, 26’933’469 were located on autosomes, and 1’263’597 were assigned to the Z chromosome (corresponding to Super-scaffold 13 and 42 in the reference assembly).

The full set of 28’197’066 high-confidence variants was phased in two successive steps. In the first step, local phasing was performed using WhatsHap v1.4 ^80^, which uses sequencing reads information to phase variants observed together on the same read or read pair, and incorporates pedigree data when available. Phasing was conducted independently for each individual. For individuals belonging to parent-offspring trios, pedigree-aware phasing was applied using WhatsHap v1.4 ^80^ pedigree mode. In cases where a parent was present in multiple trios (i.e., had more than one offspring in the dataset), the parent was phased together with the offspring that had the highest sequencing depth. Additionally, we incorporated the mean recombination rate of the species ^44^, by using the *--recombrate 2* option during phasing. After phasing, we applied a second round of variant filtering to minimize bias introduced by relatedness among individuals. Specifically, we retained only biallelic SNPs with minor allele count (MAC) > 5, and missing data < 5%, based on a subset of 187 unrelated individuals, defined as having pairwise kinship coefficients < 0.03125 (see ^44^ for kinship estimation details). Following this filtering step, the final dataset comprised 10’451’268 SNPs, of which 10’115’035 were located on autosomes and 336’233 on the Z chromosome.

The set of 10’115’035 autosomal variants was further phased using SHAPEIT v4.1.2 ^81^. This algorithm extends local phasing by incorporating population-level information using a model based on coalescence and recombination, enabling both statistical phasing of haplotypes and imputation of missing genotypes. To minimize potential bias due to relatedness, phasing was performed in two stages: 187 unrelated individuals were phased together to form a reference haplotype panel. 315 family members were then phased individually against this unrelated set, avoiding the inclusion of family structure in the phasing panel and maintaining statistical independence. SHAPEIT was run according to the authors’ recommendations for increased phasing accuracy. Specifically, the number of conditioning hapolotypes in the PBWT (Positioning Burrows-Wheeler Transform) was set to 8. The Markov Chain Monte Carlo (MCMC) parameters were set as follows: 10 burn-in iterations, 5 pruning iterations, each separated by 1 additional burn-in iteration, and 10 main iterations for haplotype sampling.

The set of 336’233 SNPs located on the Z chromosome was phased independently to account for the haplo-diploid nature of this chromosome in birds (males: ZZ, females: ZW). For variants located within the pseudo-autosomal region (PAR) – as defined in ^44^, the phasing procedure followed the same protocol as for autosomal variants, since all individuals are diploid in this region. For non-PAR regions, phasing was handled differently by sex: diploid males (ZZ) were phased using the same two-step procedure applied to autosomes: first using WhatsHap ^80^ for read-based phasing, followed by SHAPEIT v4.1.2 ^81^ for statistical phasing. Haploid females (ZW) possess a single Z chromosome; thus, any heterozygous calls (e.g., due to sequencing or genotyping error) were recoded as haploid genotypes by assigning the most frequent allele observed across individuals at that site. After sex-specific phasing, the male and female datasets were merged, and female genotypes were recoded as fully homozygous diploid for downstream analyses, ensuring compatibility with software that requires diploid input. This yielded a final dataset comprising 10’451’268 phased SNPs, including both autosomal and sex-linked variants.

To evaluate phasing quality on autosomes, we used the switch error-rate (SER) metric ^82^. A switch error occurs when a heterozygous site is phased differently between two phase sets for the same individual. For each individual, we compared local phasing inferred either form a read-based or the trio approach (both using WhatsHap ^80^ as described previously) with statistical phasing produced by SHAPEIT ^81^. In this comparison, SHAPEIT phasing was performed without using read-based or trio information for the target individual, using the same version and parameters detailed in the previous section. This method assumes that short phasing blocks generated by WhatsHap are accurate; thus, any switch in these blocks when compared to SHAPEIT phasing is considered an error. Switch error rates were calculated using the switchError tool (available at https://github.com/SPG-group/switchError). Our results show that variants present in the reference panel had a low switch error rate, with a mean error SER of 1.83% (Fig. S12).

We also evaluated the ability of the reference panel to capture both SNP diversity and the arrangement of these SNPs into haplotypes within the population. To assess SNP diversity, we downsampled the number of individuals and estimated the number of unique SNPs present in each subset. We observed that the number of SNPs plateaued after approximately 200 individuals, indicating that the reference panel effectively captures the majority of common SNPs found in the Western Palearctic population (Fig. S10). To assess haplotypic diversity, we applied a similar downsampling approach and found that the panel also captures fine-scale haplotype structure (Fig. S13).

#### Low coverage: DNA extraction, library preparation, and sequencing

We sequenced 2,778 owls from a pedigreed population, with samples collected between 1986 and 2020. Individuals were prioritized based on the availability of complete family information and phenotypic data. To assess sequencing performance, we included 32 individuals previously sequenced at high coverage as part of the reference panel (see above). Of these, 10 individuals were sequenced three times at low coverage, once per flow cell, to assess potential batch effects (see below for details). These are referred to as *triplicates*. The remaining 22 individuals, each sequenced once at low coverage, are referred to as *duplicates* and were used to evaluate the accuracy of low-coverage genotyping by comparison with their high-coverage data (see next paragraphs).

Genomic DNA was extracted from blood samples using the DNeasy Tissue Kit (Qiagen, Switzerland) and the Biosprint robot 96 (Qiagen, Switzerland), then stored at-20°C. High quality DNA (without degradation) was quantified using Quant-it PicoGreen dsDNA Assay kit (Thermo Scientific, Switzerland) and diluted in 10 mM Tris-HCl to 1.5 to 2.5 ng/μl. We randomized the position of the 2,820 DNA samples (2,778 owls + 20 triplicates + 22 duplicates) across 30 distinct 96-well plates. Each plate included two empty wells for contamination control. DNA concentration was quantified using the Quant-it PicoGreen dsDNA Assay kit (Thermo Scientific, Switzerland), and samples were diluted in 10 mM Tris-HCl to a final concentration of 1.5 to 2.5 ng/μl. Libraries were prepared using the plexWell 96 kit (SeqWell, USA) and sequenced across three lanes (10 plates per lane) on the Illumina NovaSeq 6000 platform at the Genomic Technologies Facility (GTF) of the University of Lausanne.

#### Low coverage: genotyping, phasing, imputation, and validation on autosomes

Raw sequencing reads were trimmed using Trimmomatic v.0.36 ^75^ and aligned to the barn owl reference genome (NCBI RefSeq assembly: GCF_018691265.1) ^73^ with BWA-MEM v.0.7.15 ^76^. Read alignment was followed by quality control using Qualimap v2.2.1 ^83^, which was used to estimate mean sequencing coverage across the autosomes and the Z chromosome. Across all individuals, the average genome-wide was 1.95X, with individual coverage ranging from 0.2X to 4.15X (fig S14).

We estimated genotype likelihoods for low-coverage individuals at each variant position present in the reference panel using BCFTools (*mpileup* and *call* methods), with the *-T* and *-C* options enabled ^84^. Phasing and imputation of these low-coverage samples were performed using GLIMPSE v1.1.1 ^85^. GLIMPSE leverages haplotypes from high-coverage reference panel to impute and phase genotypes in low-coverage samples. We used the 502 high-coverage individuals described earlier as the reference panel. We selected version 1.1.1 of GLIMPSE, as it is better suited for small reference panels according to the official documentation. The GLIMPSE pipeline consists of four steps: Chromosomes were divided into overlapping chunks for efficient parallel processing using the GLIMPSE_chunk tool. We applied default parameters (*--window-size 2,000,000* and *--buffer-size 200,000*). Each genomic chunk was phased and imputed using the *GLIMPSE_phase* method. This step iteratively refines genotypes likelihoods and phase information for each individual independently. In each iteration, GLIMPSE phases the low-coverage individual using haplotypes from both the high-coverage reference panel and other imputed low-coverage samples. It identifies the closest haplotypes, those sharing the highest number of identical-by-descent (IBD) segments and uses them for genotype imputation. Within each diploid sample, haplotypes are imputed separately. A new iteration begins once imputation is complete. We performed this step with an increased number of iterations (*--burnin 100* and *--main 15*) and set the effective population size parameter to (*--ne 10000*). If a recombination map was available for the given chunk ^44^, it was included using the *--map* option. All samples were considered diploid for autosomal genotyping. Once phasing and imputation were completed for each chunk, results were merged using the *GLIMPSE_ligate* tool to ensure consistent phase across the genome. This step was run with default parameters The final step involved identifying the most likely haplotype configurations based on posterior genotype likelihoods and phase probabilities, using the GLIMSE_sample method with the *--solve* flag enabled.

Validation of the imputation approach was performed using the 32 individuals that were sequenced at both high and low coverage. As previously mentioned, GLIMPSE selects haplotypes sharing the highest fraction of identical-by-descent (IBD) segments with the target sample for both phasing and imputation. Including the same individual in both the reference panel and the imputed (target) set would therefore lead to artificially accuracy due to circularity. To avoid this bias, we ran the GLIMPSE pipeline five times: We used the full reference panel of 502 individuals and the 2,768 unique low-coverage samples (excluding all duplicates and triplicates). This run produced the *main* dataset used throughout the study. The 22 duplicate individuals (those sequenced at both high and low coverage) were removed from the reference panel and added to the low-coverage target set. This resulted in a reference panel of 480 individuals and 2,790 low-coverage samples. This setup was used specifically to validate imputation accuracy without self-reference bias. Since GLIMPSE also samples haplotypes from other low-coverage individuals, we further ensured independence by imputing each of the three low-coverage replicates of the 10 *triplicate* individuals separately. In each run (denoted a, b, and c), the triplicate being imputed was excluded from the rest of the low-coverage data. These runs used as reference panel of 492 individuals (502 minus the 10 triplicates) and 2,778 low-coverage samples. To ensure fair evaluation, we excluded). Because 11 individuals from the validation set that had intermediate sequencing depth in the high-coverage dataset (mean ≈11x), as their coverage could bias the accuracy assessment (see ^86^ for details). Using the remaining n=21 individuals sequenced at high coverage (mean≈30x), we estimated the overall imputation accuracy as r^2^ = 0.968 across all variants. For further methodological details on imputation and validation procedures, refer to^86^.

### Identification of genomic regions associated with plumage coloration and spottiness

#### GRM and kinship matrix

We estimated individual-based relatedness (β) ^87^ and inbreeding coefficients using the R package hierfstat (v0.5-11, R v4.2.2) ^88^. These metrics were calculated for all individuals in the dataset. The resulting kinship matrix was then converted into a Genetic Relationship Matrix (GRM) using the *kinship2grm()* function provided by the hierfstat package. This GRM was used in downstream association analyses to account for population structure and relatedness among individuals.

#### SNPs-based heritability of plumage coloration and spottiness

We estimated SNP-based heritability for the two traits, plumage coloration and spottiness, by fitting two independent animal models. These general linear mixed models partition the total phenotypic variance (V_P_) into three components: (1) variance explained by fixed effects, accounting for known environmental sources; (2) additive genetic variance (V_A_), estimated via the GRM, and (3) residual variance (V_R_). Animal models were implemented in R using the brms package (version 2.34, R version 4.2.2) ^58^. Sex and phenotypic stage were included as fixed effects to control for their influence on trait variation. The GRM, previously described, was incorporated as a random effect to estimate V_A_. Models were run using 8 independent Markov Chain Monte Carlo (MCMC) chains, each with 10,000 iterations and burn-in of 2,000 iterations. Narrow-sense heritability (h^2^) was calculated as h^2^=V_A_/V_P_. We also report the 95% credible intervals associated with each h^2^ estimate. Results for both traits are presented in Supplementary Table S9 and S10.

#### Genome-Wide Association Study (GWAS)

To identify genomic variants associated with plumage coloration and spottiness in the Swiss barn owl population, we performed genome-wide association analyses using the *association.test()* function from the *gaston* package (v1.6, R v4.2.2) ^89^. A linear mixed model (LMM) was fitted using the Average Information Restricted Maximum Likelihood (AI-REML) algorithm, as implemented in the same package ^89^. Both traits were treated as continuous variables. Sex and phenotyping stage (juvenile or adult) were included as fixed covariates, while the Genetic Relationship Matrix (GRM) was included as a random effect to account for population structure, familial relationships, and cryptic relatedness. To avoid proximal contamination and increase statistical power, we applied a Leave-One-Chromosome-Out (LOCO) approach. For each chromosome, we constructed a GRM excluding SNPs from that chromosome, and performed the GWAS on the focal chromosome using this modified GRM. This approach ensures that associations are not inflated due to shared signal between the tested SNPs and the GRM ^90^. Mixed-model association methods incorporating LOCO have been shown to outperform traditional GWAS approaches in terms of power and control for confounding ^91^.

For every GWAS, we used the score statistics to assess the strength of the association between individual SNP and the phenotype of interest. The significance threshold (α = 0.05) was adjusted using a Bonferroni correction to account for multiple testing ^92^. Additionally, we visually assessed deviations from the null distribution using quantile-quantile (QQ) plots, focusing on SNPs deviating from the expected 1:1 line as a qualitative check for inflation or strong associations.

To maximize power for detecting loci associated with plumage coloration and spottiness, we conducted six independent GWAS, three for each trait category.

For the coloration, we first ran a GWAS using the mean ventral plumage coloration as the phenotype. However, since the coloration of the belly and breast are not perfectly correlated (Fig. S3), we conducted two additional GWAS, using belly coloration and breast coloration as independent traits. This allowed us to identify loci potentially contributing to regional variation in coloration patterns.

For the spottiness, we began with a GWAS on total spottiness as a composite trait. Given that this phenotype results from the combination of both spot number and spot diameter, we stratified the dataset to explore the genetic basis of each component separately. This involved running two additional GWAS: (i) one using the number of dots, and (ii) one using the mean spot diameter as the phenotype.

### Genotyping and gene expression of the Z chromosome region

#### Feather and blood sampling and genotyping

For gene expression and allelic discrimination, we collected two to three developing breast feathers and blood samples from 28 male and 17 female nestlings born in 2023 (n= 21) and 2024 (n = 24). We collected growing feathers from nestlings at a similar stage of development (mean age ± SD: 29 days ± 1.8). At this age, the feathers start to develop the typical black spots on the apical part of the feather and the white to reddish pheomelanin coloration. Upon collection, feathers and blood were immediately frozen in liquid nitrogen in the field and stored at-80° C until molecular analyses. For every individual, genomic DNA was extracted using the Blood and Tissue kit (Qiagen, Switzerland), sexed (as described in ^67^) and genotyped for *MC1R* (as described in ^57^). Out of the 45 individuals used for the gene expression analysis, The *MC1R* genotypes are the following: 38 *VV* (GG genotype, homozygous MC1R_white_), 6 *VI* (AG genotype), 1 male *II* (AA genotype, homozygous MC1R_red_). Because of this unequal frequencies, *MC1R* genotypes were not considered for the analysis.

#### Allelic discrimination assay

To validate associations found on the Z chromosome, we performed an allelic discrimination assay targeting a SNP located on Super-Scaffold 42 at position 29,811,381. This SNP was selected because it showed strong linkage disequilibrium with the top associated SNPs for both plumage coloration (r² = 0.97) and spottiness (r² = 0.98) and was itself significantly associated with both traits (p < 6.95e-31 for coloration; p < 2.70e-122 for spottiness). The following primers and probes were used in the assay: Forward primer ChZ29811429Fw: 5’-TATCTTTGGGCTTGACTGGT-3’; Reverse primer: ChZ29811308Rev: 5’-AAACACCCAAGAAATAGCAAT-3’; Probe 1 (FAM-labeled): ChZ29811381FamMGBQ530: 5’-ACTCTCACTTTGTTGCTCTCTCCT-3’; Label: 5’ FAM fluorophore; 3’ MGB quencher; Probe 2 (HEX-labeled): ChZ29811381HexMGBQ530: 5’-ACTCTCACTTTGCTGCTCTCTCCT-3’ Label: 5’ HEX fluorophore; 3’ MGB quencher.

To enhance signal quality, an initial PCR pre-amplification step was performed using the following thermal cycling conditions: 95°C for 5 minutes (Initial denaturation) followed by 30 cycles at 95°C for 30 seconds, 56°C for 30 seconds, 72°C for 30 seconds, followed with 72°C for 10 minutes (final extension). The reaction mix (10 μL total volume) contained: 200 μM dNTPs, 1 mM MgCl2, 500 nM of each primer, 1 x GoTaq buffer, 0.5 U/μl GoTaq polymerase (Promega, Switzerland). Following PCR, the products were diluted 1:1,000,000, and 2 μL if the diluted product was added to a 20 μL qPCR reaction using Takyon MasterMix (1x) (Eurogentec, Belgium), with 300 nM of each primer and 300 nM of each probe.

qPCR and allelic discrimination were performed on a CFX96 Real-Time PCR Detection System (BIO-RAD, Switzerland) using the standard protocol. The assay was conducted on 184 owls previously sequenced at low coverage. We observed a very high concordance between the genotypes inferred from allelic discrimination and those obtained via imputation from low-coverage whole-genome sequencing, with r^2^ = 0.986. Only one individual showed a discrepancy; further inspection revealed that this sample had no sequencing coverage at the locus, suggesting the discordant imputed genotype was due to a miscall during imputation rather than an error in the allelic assay.

The allelic discrimination for the region on the Z chromosome (SNP 29,811,381) was then used to genotype the 45 individuals used for RNA extraction and gene expression. The genotypes at the Z locus were for males 5 *CC*, 13 *CT*, 10 *TT*, and for females 5 *C* and 12 *T*.

#### Total RNA extraction and cDNA preparation

Total RNA was extracted from the developing breast feathers of the 45 nestlings used in the gene expression study. For each individual, one to two growing feathers were used. At the time of RNA extraction, a photograph of the basal part of each feather was taken on dry ice to visually document the stage of black spot development. Feathers were ground in liquid nitrogen using a sterile pestle and then resuspended in Qiazol Lysis Reagent. Total RNA was extracted using the miRNeasy Mini Kit (Quiagen, Switzerland) following the manufacturer’s protocol, with the following modifications: RNase-Free DNase treatment (Quiagen) was included during the extraction to remove genomic DNA; Buffer RWT was prepared using isopropanol instead of ethanol, as recommended for feather samples. Total RNA was eluded in 30 μL of RNase-free H_2_O. RNA concentration was quantified using the Qubit fluorometer (Life Technologies, Switzerland) with the RNA Broad Range Assay Kit, and RNA integrity was assessed using the Fragment Analyzer (Advanced Analytical, Labgene, Switzerland). Only samples with high RNA quality (RQN > 7.7) were retained for downstream gene expression analyses. For each individual, 200 ng of total RNA was reverse-transcribed in a total volume of 20 μL using the LunaScript RT SuperMix (New England Biolabs; BioConcept, Switzerland), under the following thermal conditions: 25°C for 2 minutes; 60°C for 10 minutes; 95°C for 1 minute (inactivation). The resulting cDNA was precipitated using 1 μL of glycogen (20 ng/μL), 5 M ammonium acetate, and ethanol, and resuspended in 20 μL of 10 mM Tris-HCl (pH 8.0) and 0.1 mM EDTA. Because of the low expression levels of some target genes, 5 μL of the precipitated cDNA were preamplified with the complete primer mix used for the qPCRs. Preamplification was performed for 14 cycles using reagents from Life technologies (Thermo-Fisher Scientific, Switzerland). The resulting product was diluted 10-fold in 10 mM Tris–HCl (pH 8.0) and 0.1 mM EDTA, and stored at −20 °C until qPCR. Preamplification was validated not to bias relative expression levels for the majority of genes. However, one reference gene (HPRT1) showed altered expression post-preamplification and was therefore excluded from all downstream analyses.

#### Gene expression analysis

To investigate the molecular basis of plumage coloration and spottiness, we quantified the expression of genes located within and adjacent to the Z chromosome region (Super Scaffold 42) previously identified as being strongly associated with both traits. Specifically, we measured the expression of 12 genes spanning the genomic interval between 29,210,880 and 30,121,259 using reverse transcription quantitative PCR (RT-qPCR). The targeted genes include: *AGGF1* (Angiogenic Factor With G-Patch And FHA Domains 1); *AP3B1* (Adaptor Related Protein Complex 3 Subunit Beta 1; *CRHBP (LOC10436293)* (Corticotropin Releasing Hormone Binding Protein); *F2R* (Coagulation Factor II Thrombin Receptor); *IQGAP2* (IQ Motif Containing GTPase Activating Protein 2); *LOC116965528* (uncharacterized non-coding RNA); *LOC122154642* (uncharacterized non-coding RNA; *PDE8B* (Phosphodiesterase 8B); *SCAMP1* (Secretory Carrier Membrane Protein 1); *SNORA47* (Small Nucleolar RNA, H/ACA Box 47); *TBCA* (Tubulin Folding Cofactor A); *WDR41* (WD Repeat Domain 41). Among these, *PDE8B* and *LOC116965528* are located closest to the SNPs showing the highest associations with spottiness and coloration. Given that multiple splice variants have been annotated for many of these genes in the T. alba_DEE_v4.0, barn owl genome assembly (NCBI RefSeq assembly: GCF_018691265.1) ^73^, we designed and tested multiple primer and probe sets per gene to evaluate potential transcript-specific expression. All primers and probes used in this study are listed in Table S17. Four additional genes located in the same genomic region could not be amplified despite multiple attempts and were therefore excluded from the expression analyses. To normalize gene expression in the RT-qPCR assays, we evaluated the expression of four commonly used reference genes: *EEF1A* (*Elongation factor 1A*)*; GAPDH* (Glyceraldehyde-3-Phosphate Dehydrogenase); *HPRT1*(Hypoxanthine Phosphoribosyltransferase 1); *TBP* (TATA-Box Binding Protein). However, due to altered expression following preamplification, *HPRT1* was excluded from all analyses. In addition to the genes located on the Z chromosome, we also quantified the expression of four candidate pigmentation genes: *MC1R* (Melanocortin 1 Receptor); *GPR143* (G Protein-coupled receptor 143); *TYR* (Tyrosinase); *TYRP1* (Tyrosinase-related protein 1). Primers and probes for these pigmentation genes were previously described ^48,93^.

Quantitative PCR (qPCR) conditions were optimized for each primer and probe set by testing various concentrations using serial dilutions of plasmids or PCR-purified products amplicons of the target gene regions. The goal was to achieve PCR efficiencies between 95% and 105%, in line with MIQE guidelines. Most primer-probe combinations fell within this range (see Table S17). One exception was the IQGAP2_T102 primer-probe set, which showed a slightly elevated efficiency of 107 %, but was retained for downstream analyses given the acceptable range for relative quantification (see below). qPCR reactionsqPCRs were performed on a CFX96 Real-Time PCR Detection System Bio-Rad (BIoRad, Switzerland). Each reaction contained 2 ul of diluted, pre-amplified cDNAs, run in technical duplicates, using the QuantiTect Probe PCR Kit (Qiagen, Switzerland) in a final reaction volume of 20 μL. To ensure technical accuracy reactions were repeated if the Ct difference between duplicates exceeded 0.25 cycles. To control for plate-to-plate variability and potential pre-amplification biases, three pooled pre-amplified cDNA controls (from different individuals) were included on every qPCR plate.

We set up qPCR conditions with different concentrations of primers and probes with serial dilutions of plasmids or PCR purified products of the amplified gene region to achieve PCR efficiency between 95% and 105% (Table S17). Only for one primer pair and probe for IQGAP2_T102, we got a higher efficiency of 107 % (Table S17) while considering the relative expression calculation (see below).

To identify the most stable reference genes for qPCR normalization, we used the RefSeeker package (v1.0.4, R v4.2.2) ^94^. RefSeeker evaluates reference gene stability using four established algorithms-Normfinder, geNorm, delta-Ct, and BestKeeper-based on the raw quantification cycle (Cq) values. Among the three tested reference genes (*EEF1A*, *GAPDH*, and *TBP*), *EEF1A* and *TBP* were identified as the most stable, with final stability rankings of 1.2 and 1.4, respectively (Table S17). These two genes were therefore used for normalization in subsequent expression analyses. All raw Ct values were log transformed to calculate relative expression assuming a PCR efficiency of 2.0 (corresponding to 100% efficiency, within the acceptable 95-105% range). For IQGAP2_T102, which had a slightly higher empirically determined efficiency, a coefficient of 2.07 was applied. Normalized relative expression values were obtained using the SL1PCR package (v1.72.0 R v4.2.2) ^95^, with EEF1A and TBP serving as internal controls.

#### Gene expression analysis and statistical modeling

All qPCR-derived gene expression data were square root-transformed to improve normality of the residuals prior to statistical analyses, using R v4.4.2. Because multiple transcript variants were detected for several genes, we first assessed the correlation between their expression levels using males and females (see Figure S21-S22). Given the high correlation observed, only one transcript per gene was retained for downstream analysis. Where possible, we selected primers and probes that either (i) targeted all transcript variants, or (ii) captured the most highly expressed variant (e.g., for *IQGAP2*, see Table S17). To test whether genotype at the Z-linked locus (SNP at position 29,811,381 on Super-Scaffold 42) influenced gene expression, we fitted linear mixed-efects model using the *lme()* function from the *nlme* (v3.1-168, R v4.2.2) ^96^.

Because males (ZZ) and females (ZW) differ in their ploidy at the Z chromosome, we ran sex-specific models. Each model included gene expression as the response variable, genotype at the Z locus as a fixed effect (additive coding), brood of origin as a random effect to account for shared environmental and genetic background. For each gene-sex combination, we compared two models: one with and one without genotype as a fixed effect. Model comparison was performed using likelihood ratio tests (ANOVA). The resulting p-values were corrected for multiple testing using the Bonferroni method ^92^. Only corrected p-values below the significant threshold (α=0.05) were considered evidence that genotype significantly affected gene expression.

For genes showing a significant association between genotype and gene expression, we further investigated whether variation in gene expression was associated with individual phenotypic traits. To do so, we fitted linear mixed models using the *lme()* function from the *nlme* package (v3.1-168, R v4.2.2) ^96^. As feathers were collected from the breast of the fledglings, we focused this analysis on plumage coloration and spottiness traits measured on the breast only. For each gene, we built two separate models: a baseline model including only sex as fixed effect; a full model including both the sex and gene expression levels as fixed effects. In both models, brood of origin was included as a random effect to account for non-independence among siblings.

### Genetic architecture of plumage coloration and spottiness

#### Variance partitioning among QTLs for color and spottiness

To quantify the contribution of the genomic loci identified in the GWASs and to evaluate the sex-specific genetic architecture of plumage coloration and spottiness, we fitted Bayesian generalized linear models using the R package brms (v2.0.22, R v4.4.2) ^58^. Given the well-documented sexual dimorphism in the species, we constructed independent models for males and females for each trait. For each trait/sex combination, we considered multiple model parameterizations to explore the nature of genotype-phenotype relationships. In one set of models, genotypes were encoded as dosage values ranging from 0 to 2, assuming additive effects of alleles. In an alternative set, genotypes were treated as categorical factors, allowing for non-additive and dominance effects at each locus.

To identify the most appropriate model for each trait and sex, we compared the prediction accuracy of all fitted models using the widely applicable information criterion (WAIC). WAIC is a Bayesian model selection tool that evaluate the log-likelihood averaged across posterior simulations, providing a robust metric for model comparison that outperforms AIC and DIC in complex hierarchical models ^97^. This approach is implemented in the brms function *loo_compare*(). Model comparison and selection were implemented using the loo_compare() functions in the brms package. This approach allowed us to identify the genetic model best explaining trait variance across sexes and traits, offering insights into sex-specific dominance or additive genetic effects at key QTLs.

In the models for plumage coloration, we included as fixed effects the genotype at the three loci identified as significantly associated with the trait in the GWAS analyses: the SNP on Super-Scaffold 26, position 22,522,039 (hereafter referred to as genoMC1R); the SNP on Super-scaffold 42, position 29,808,233 (genoZcol); the SNP on Super-scaffold 6, position 27,439,651 (genoCORIN)). We also included the developmental stage at phenotyping (fledglings versus adults) as a fixed factor, to control for age-related variation in coloration. To account for relatedness among individuals and estimate the additive genetic variance (V_A_), the genomic relatedness matrix (GRM) calculated from autosomal SNPs was included in the models as a random effect. To investigate potential epistatic interactions between loci, we additionally fitted two extended models including the interaction term between genoMC1R and genoZcol. All models assumed a Gaussian distribution for the coloration trait. Bayesian inference was performed using 8 independent Markov chains, each run for 10,000 iterations, with a burn-in of 2,000 iterations. We set the *adapt_delta* parameter to 0.9 to ensure proper convergence.

For the analysis of spottiness, we included as fixed predictors the genotype of individuals at the two loci significantly associated with the trait in the GWAS: the SNP on Super-scaffold 42, position 29,812,087 (hereafter referred as genoZspot); the SNP on Super-scaffold 26, position 22,522,039 (genoMC1R). We also included the developmental stage at which the phenotype was measured (fledgling or adult) as a fixed effect. To account for genetic relatedness, we included the genomic relatedness matrix (GRM), estimated across the entire genome, as random effect in the models, allowing us to estimate *V_A_*. Because the distribution of the spottiness trait was zero-inflated, with an over-representation of individuals showing no spots, we modeled the phenotype using a hurdle-gamma distribution. In this framework, the presence or absence of spots was modeled as the binary hurdle component, while the degree of spottiness among spotted individuals was modeled with a gamma distribution. All models were run using 8 independent Markov chains, each for 10,000 iterations, with a burn-in of 2,000 iterations. The *adapt_delta* parameter was set to 0.9 to ensure robust convergence. For the best-fitting models, we extracted the total variance explained using the *bayes_R2()* function from the *brms* package ^58^.

### Allele frequency changes, fitness estimates, and drivers of evolution

#### Temporal evolution of allele frequencies

To assess whether the allele frequencies at the loci identified by the GWAS for coloration and spottiness have changed over time, we estimated allele frequencies for each year based on the adults alive in that year. To account for variation in sample sizes, particularly in earlier years with limited data, we computed 95% confidence intervals around each annual allele frequency estimate using the Clopper–Pearson exact binomial method ^98^.

To evaluate whether allele frequencies have changed systematically over time, we fitted linear models with allele frequency as the response variable and year as the predictor. To avoid potential bias from years with high uncertainty in frequency due to small sample sizes, we restricted this analysis to the period between 1995 and 2022.

#### Estimation of Effective population size

We estimated the effective population size (*Ne*) from the whole-genome resequencing data using the standard formula *π*/(4\**Mu*), where *π* is the nucleotide diversity and *μ* the mutation rate per site per generation [REF]. For each year, *π* was calculated using only for the adult individuals alive during that year. To ensure consistency with the variant calling pipeline, the total sequence length (L) used for *π* estimation matched the length of the reference genome after mapping (L = 1,131,781,302 bp). Nucleotide diversity (*π*) was computed using the *pi.dosage()* function from the hierfstat (v0.5-11, R v4.2.2) ^88^. Because the mutation rate in barn owls is currently unknown, we adopted the recently estimated mutation rate for the closely related Snowy owl (*Bubo scandiacus*) (Mu = 1.93*10^-9^) ^99^. To obtain 95% confidence intervals around annual Ne estimates, we performed bootstrap resampling of non-overlapping 1Mb segments along the genome and repeating this process 100 times per year.

#### Simulation of allele frequency through time

To assess whether the observed allele frequency changes could be explained solely by genetic drift, we simulated allele frequency dynamics under a neutral scenario. This approach allowed us to test whether selection may have also played a role in shaping the temporal trajectory of alleles associated with coloration and spottiness. We modeled drift in a population with overlapping generations, reflecting the demography of barn owls, which have an average lifespan of approximately three years^100^. Accordingly, in our simulations, one-third of the population was renewed each year, while the remaining two-thirds persisted from the previous generation. Simulations began with the initial allele frequencies observed in the population over the first four years of the study: *MC1R_red_* 0.047; Z_spotted_ 0.685 (based on n = 32 individuals). For each locus, we ran 1000 replicates assuming a constant population size of 400 individuals, a conservative estimate consistent with the lowest census sizes of the Swiss barn owl population during the study period ^63^. Given that *MC1R* is located on an autosome, simulation assumed diploidy across all individuals. The Z locus is located on the Z chromosome, which is hemizygous in females (ZW) and homozygous in males (ZZ). We therefore used a sex-aware simulation approach, where the effective number of alleles per individual differed by sex (1 for females, 2 for males), and sexes were sampled at a 1:1 ratio. To mimic the empirical data structure, for each simulated year we subsampled the same number of individuals as were observed alive in the actual dataset for that year. This allowed us to account for sampling variance and generate realistic confidence intervals around the neutral expectation.

#### Disentangling drift and selection on major QTLs

To evaluate whether the allele frequency changes observed at key QTLs were consistent with neutral expectations or indicative of selection, we used the approxWF framework ^101^. This method compares empirical allele frequency trajectories to those expected under a Wright–Fisher diffusion process and is particularly well suited for time-series population genetic data. It allows explicit estimation of selection coefficients while accounting for stochastic fluctuations due to genetic drift. For each focal locus, we fitted two models: (i) a neutral model (drift only), and (ii) a selection incorporating a selection coefficient (s) and, optionally, a dominance coefficient (h). Each Markov Chain Montel Carlo (MCMC) chain was run for 1,010,000 iterations, with the first 10,000 iterations discarded as burn-in, and sampling conducted every 10 steps. For the Z-linked locus, we used the default MCMC parameters. For MC1R, however, the default settings led to low acceptance rates and unstable chains, likely due to lower starting allele frequency. To improve convergence and stability, we reduced the proposal step sizes by half, using logN_step=0.025 for the log population size, s_step=0.005 for the selection coefficient (s), and h_step=0.005 for the dominance coefficient (h).

For each locus, we first fitted a Wright-Fisher model incorporating both genetic drift and selection, and compared the log-likelihoods and the proportion of MCMC iterations supporting selection versus neutrality. A higher posterior support for selection and an improved model likelihood was interpreted as evidence that selection better explains the observed allele frequency trajectory. We then fitted a selection-only model and assessed whether the 95% credible interval of the estimated selection coefficient excluded zero. A locus was considered under directional selection if it met the following three criteria: (i) the selection model had a higher log-likelihood than the drift-only model; (ii) the 95% credible interval did not include zero; and (iii) a substantial proportion of MCMC iterations supported the presence of selection. Loci that failed to meet these criteria, particularly those with overlapping credible intervals including zero and low support for selection, were interpreted as being consistent with neutral drift. To provide further insights into the evolutionary dynamics, effective population size and dominance coefficients were jointly inferred from the MCMC chains.

Finally, we used the simulation and dominance coefficients estimated from the selection-only models to simulate allele frequency trajectories under selection. Simulations followed the scheme outlined in the *Simulation of Allele Frequency Through Time* section. Genotype fitness was parameterized as 1, 1-hs, and 1-s for autosomal loci (MC1R), and as 1 and 1-s for hemizygous females at the Z-linked locus.

## Author contributions

T.C. and J.G. were responsible for the conceptualization of the study, with contributions from A.T., E.L., A.L.D. and A.R. B.A. and A.R. supervised sample acquisition and individual phenotyping over the long-term study period. A.L.D. supervised sample preparation for sequencing and carried out the gene expression experiment, with contributions from C.S. A.T. assembled the reference panel with contributions from T.C. and E.L. E.L. performed genotyping and imputation of the low-coverage samples, with contributions from T.C., A.T., and S.S.C. T.C. conducted the formal analysis, with assistance from P.B. for phenotypic analyses, A.H. for the quantitative genetic approach, and A.L.D. for the gene expression analyses. A.T., E.L., and J.G. contributed to the analyses throughout the process.

A.R. and J.G. were responsible for funding acquisition. T.C. wrote the original draft, with contributions from A.T., E.L., A.L.D., and J.G. All authors read, provided feedback on, and approved the final version of the paper.

## Acknowledgement

We thank all individuals who contributed to this long-term project, including field assistants, former PhD students, postdoctoral researchers, and collaborators (full list in Supplementary Information). We are also grateful to the Genomic Technologies Facility (GTF), Lausanne, for sequencing support and to Olivier Delaneau for insightful discussions.

## Data availability

The data underlying this article are available in the GenBank Sequence Read Archive Database at https://www.ncbi.nlm.nih.gov/sra. High coverage sequences used for the reference panel can be accessed within BioProject PRJNA700797, PRJNA727915, PRJNA727977, PRJNA774943, PRJNA925445 and PRJNA1172395. Low coverage sequences will be accessible upon publication. Code used to prepare the genomic data can be found at: https://github.com/cumtr/Cumer_et_al_EvolutionColor_Analyses

Code used for downstream analyses can be found at: https://github.com/cumtr/Cumer_et_al_EvolutionColor_Analyses

