## Supplementary material for "Genomic bases of short-term evolution in the wild revealed by long-term monitoring and population-scale sequencing": TableS17_primersRNA.docx

**Supplementary Table** : Sequences and efficiency of the probe and primer pairs used to quantify the gene present on the Z chromosome

| Primer name | Primer sequence | Gene name | Start position of the gene | | Stop position of the gene | | Accession #^1^ of the transcript | | qPCR efficiency (%) | | R^2^ |
| --- | --- | --- | --- | --- | --- | --- | --- | --- | --- | --- | --- |
| AGGF1_1231_F | GGATCACTTTATATTATCACAGCGG | *AGGF1* | 29835465 | | 29860070 | | XM_042804694.1 | | 105 | | 1 |
| AGGF1_1287_FamQ1 | TGTTGGACACACACTTCAGATTCCGGA |  |  | |  | |  | |  | |  |
| AGGF1_1373_R | ACATAACTTTGCAAATCATGGTCA |  |  | |  | |  | |  | |  |
| AP3B1_all_253F | TGTGAAGAATGTTGCCAGTAA | *AP3B1* | 29300483 | | 29461212 | | XM_033009138.2 | | 98 | | 0.992 |
| AP3B1_all_348FamQ1 | TGTCCATCAGCACTTTCCAACGCGCTTTA |  |  | |  | | XM_042804700.1 | |  | |  |
| AP3B1_all_402R | GCACGGATTAACTGGTTAGG |  |  | |  | | XM_042804699.1 | |  | |  |
| AP3B1_T138_785F | AAGGATGAGGTGGTGGAAG | *AP3B1* | 29300483 | | 29461212 | | XM_033009138.2 | | 99 | | 0.996 |
| AP3B1_T138_834FamQ1 | CTGACGAAGAACAGAAGGAAAATGATCGCAG |  |  | |  | | XM_042804700.1 | |  | |  |
| AP3B1_T138_893R | CTGGGTCCATAGTATAGACGT |  |  | |  | |  | |  | |  |
| AP3B1_T698-700_714F | AAGAGTGGGGTCAAGTTGTT | *AP3B1* | 29300483 | | 29461212 | | XM_042804698.1 | | 100 | | 0.998 |
| AP3B1_T698-700_739FamQ1 | TCACATGTTAACTCGATATGCACGTACACAGT |  |  | |  | |  | |  | |  |
| AP3B1_T698-700_800R | CCACCTCATCATCCTTCCAT |  |  | |  | |  | |  | |  |
| AP3B1_T699_604F | TACGGATGAGGTGGTGGAA | *AP3B1* | 29300483 | | 29461212 | | XM_042804699.1 | | 98 | | 1 |
| AP3B1_T699_834FamQ1 | CTGACGAAGAACAGAAGGAAAATGATCGCAG |  |  | |  | |  | |  | |  |
| AP3B1_T699_900R | CTGTGATCTGGGTCCATAGTAT |  |  | |  | |  | |  | |  |
| AP3B1_T700_2963F | CTTTTGCCTGTCACTATGTCA | *AP3B1* | 29300483 | | 29461212 | | XM_042804700.1 | | ND | | ND |
| AP3B1_T700_3007FamQ1 | AGCCTGCCTCTCTGCTTTATACCTGCTCA |  |  | |  | |  | |  | |  |
| AP3B1_T700_3071R | TGTCTCATTCATTCCTGCCA |  |  | |  | |  | |  | |  |
| CRHBP_290F | ACGACTTTGTGAACATTGACTG | *CRHBP / LOC104362934* | 29893179 | | 29902228 | | *XM_009969317.3* | | 100 | | 0.999 |
| CRHBP_312FamQ1 | CCAAGGGGGCGATTTCCTGAGGGTAT |  |  | |  | |  | |  | |  |
| CRHBP_359R | CCTTTGAGTATCCAGCCATCAA |  |  | |  | |  | |  | |  |
| F2R_662F | CGTCCTGTTTGTAAGCGTG | *F2R* | 29977647 | | 29986244 | | XM_042804909.1 | | 100 | | 0.998 |
| F2R_708FamQ1 | TCTGGAAACGACTGGATTTTTGGACCTCA |  |  | |  | |  | |  | |  |
| F2R_771R | TACAGTAGAAGGCAGCAGTG |  |  | |  | |  | |  | |  |
| IQGAP2_T102_976F | GTTTGAACCGTCCTGCTCTG | *IQGAP2* | 29990457 | | 30121259 | | XM_033009102.2 | | 108 | | 0.999 |
| IQGAP2_T102_1090R | CACACAATTGCAATAGGTACCATGA |  |  | |  | |  | |  | |  |
| IQGAP2_T102_995FamQ1 | AGCCTAGTTGAAGTTCTGCCGCAA |  |  | |  | |  | |  | |  |
| IQGAP2_T103_1210R | GGATCACAGTTACGAATTGCAGT | *IQGAP2* | 29990457 | | 30121259 | | XM_033009103.2 | | 98 | | 1 |
| IQGAP2_T103_846F | GAGATGTCTACGAGGAGCTGC |  |  | |  | |  | |  | |  |
| IQGAP2_T103_868FamQ2 | AACACAAGCTGAGATTCAAGGCAACATCAA |  |  | |  | |  | |  | |  |
| IQGAP2_T104_1665FamQ1 | AAAACTCTAGAGACCCTACTGTTGCCCACA | *IQGAP2* | 29990457 | | 30121259 | | XM_033009104.2 | | 104 | | 0.998 |
| IQGAP2_T104_1716R | GTCTCACATCTTGCAGCTTG |  |  | |  | |  | |  | |  |
| IQGAP2_T104_902F | ATAAACAGAATCATTGCAATTGG |  |  | |  | |  | |  | |  |
| IQGAP2_T105_1716R | GTCTCACATCTTGCAGCTTG | *IQGAP2* | 29990457 | | 30121259 | | XM_033009105.2 | | 100 | | 0.999 |
| IQGAP2_T104_1665FamQ1 | AAAACTCTAGAGACCCTACTGTTGCCCACA |  |  | |  | |  | |  | |  |
| IQGAP2_T105_883F | TCAAGGAATCATTGCAATTGG |  | 29990457 | | 30121259 | |  | |  | |  |
| LOC116965528_3309F | GATCAATGCGAGCTGTGCC | *LOC116965528* | | 29828695 | | 29833607 | | XR_004409438.2 | | 100 | |
| LOC116965528_3355FamQ1 | TGGGATTTGTCACACCAATGCTAAACCGA |  |  | |  | |  | |  | |  |
| LOC116965528_3436R | CTGCAGCTTCCTCTTTCAGC |  |  | |  | |  | |  | |  |
| LOC122154642_119FamQ1 | CAGATCCTGAGTCGACTCCCTGGAGCC | *LOC122154642* | 29650507 | | 29653003 | | XR_006166676.1 | | 102 | |  |
| LOC122154642_145R | GCTCTCCAGATTTCCTACCA |  |  | |  | |  | |  | |  |
| LOC122154642_27F | AGTCCGTTGGTGTCTGATTT |  |  | |  | |  | |  | |  |
| PDE8B_all_271F | CCACGCACATCTACTGACCA | *PDE8B* | 29'725'424 | | 29'805'725 | | XM_042804417.1 | | 100 | |  |
| PDE8B_all_301FamQ1 | TCTGTTCTTCCCCTTCTTAATGCAGGCTT |  |  | |  | | XM_042804418.1 | |  | |  |
| PDE8B_all_413R | TGAGATCGCACTTCCCCATG |  |  | |  | |  | |  | |  |
| PDE8B_T17-9_-40FamQ1 | CGGCAAGCAGGCTTCAACTACAGAAGTACA | *PDE8B* | 29'725'424 | | 29'805'725 | | XM_042804417.1 | | 100 | |  |
| PDE8B_T17-9_-63F | GAGACCCAGACCAGTAACGC |  |  | |  | |  | |  | |  |
| PDE8B_T17-9_17R | TCTTGTGTCAGTCTCATGGGC |  |  | |  | |  | |  | |  |
| PDE8B_T18_-40F | CGGCAAGGCTTCAACTACAG | *PDE8B* | 29'725'424 | | 29'805'725 | | XM_042804418.1 | | 99 | |  |
| PDE8B_T18_-6FQ1 | GGGCCCATGAGACTGACACAAGATCCA |  |  | |  | |  | |  | |  |
| PDE8B_T18_90R | CTCTATCACAAGCCCACCAGA |  |  | |  | |  | |  | |  |
| SCAMP1_229F | GCGAAGGAGCATGCCTTG | *SCAMP1* | 29210880 | | 29260692 | | XM_033009107.2 | | 98 | |  |
| SCAMP1_250FamQ1 | AGGCAGAACTGCTAAAGCGTCAAGA |  |  | |  | |  | |  | |  |
| SCAMP1_309R | GCGATCTAGTTCAGCTGCTTT |  |  | |  | |  | |  | |  |
| SNORA47_121R | GTACTGCCTTCCCTCATGTCA | *SNORA47 / LOC116965530* | 29833008 | | 29833136 | | XR_004409439.1 | | 97 | |  |
| SNORA47_1Fw | TGGAGGACTGAGAAGGTGAGA |  |  | |  | |  | |  | |  |
| SNORA47_76FamQ1 | GCAATGGCAACTCATGTTCCCCTCC |  |  | |  | |  | |  | |  |
| TBCA_121F | GACCAGCGCTTGAGGAAG | *TBCA* | 29594642 | | 29626100 | | XM_042804862.1 | | 100 | |  |
| TBCA_139FamQ1 | ATCAAGATCAAGGCTGGCGTCGTCA |  |  | |  | |  | |  | |  |
| TBCA_259R | TGTAATCATCGCATGCTTCAG |  |  | |  | |  | |  | |  |
| WDR41_all_617F | TCAAAGACTGGATGTATGGTTG | *WDR41* | 29700513 | | 29744781 | | XM_033009046.2 | | 100 | |  |
| WDR41_all_641FamQ1 | TGGAGGGAGTGATCTATGTGTTTGGAACCA |  |  | |  | | XM_042804420.1 | |  | |  |
| WDR41_all_732R | CAACCAAAGCACTGATACCTG |  |  | |  | | XM_042804421.1 | |  | |  |
| WDR41_T1_1308F | GGAAGGGCAAACAAACAGGC | *WDR41* | 29700513 | | 29744781 | | XM_033009046.2 | | 100 | |  |
| WDR41_T1_1375FamQ1 | TGGAGCTGGTTGGTGATTTGATTGGTCA |  |  | |  | |  | |  | |  |
| WDR41_T1_1458R | GTAGCCAGACCCAGTTCACC |  |  | |  | |  | |  | |  |
| WDR41_T2_1370F | TCTTGGAGCTGGTTGGTGAT | *WDR41* | 29700513 | | 29744781 | | XM_042804420.1 | | ND | |  |
| WDR41_T2_1417_FamQ1 | AGAAGCAAGGGGAAGAGCAGGTGGAGTG |  |  | |  | |  | |  | |  |
| WDR41_T2_1453R | GCAAAACTACTCCAGAACCACA |  |  | |  | |  | |  | |  |
| WDR41_T3_1397F | TGGTCATTCATCAGCTGTGCA | *WDR41* | 29700513 | | 29744781 | | XM_042804421.1 | | 100 | |  |
| WDR41_T3_1439FamQ1 | AAAGAACAGAGACCCCAGCAAGCTT |  |  | |  | |  | |  | |  |
| WDR41_T3_1498R | ACATGGTGGTTATCTCCTTTTGA |  |  | |  | |  | |  | |  |

1: # = number

ND= not detected
