## Supplementary material for "Genomic bases of short-term evolution in the wild revealed by long-term monitoring and population-scale sequencing": TytoEvolutionColor_v1_BioRxiv_SupMat.docx

Supplementary Materials

### Phenotypic changes in the two melanic traits

#### Color and Spottiness variation in the Swiss barn owl

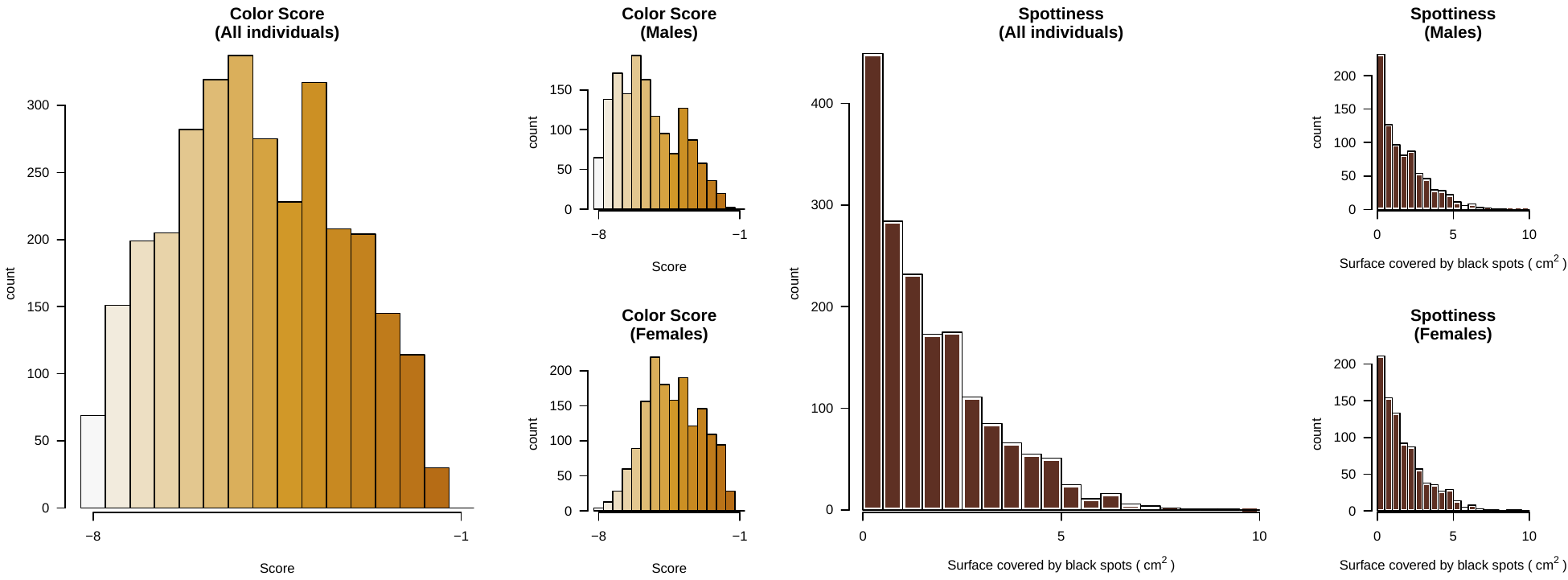

**Figure S1** – Distribution of the color (left) and spottiness (right) in all individuals and within each sex (top panel for males, bottom panel for females).

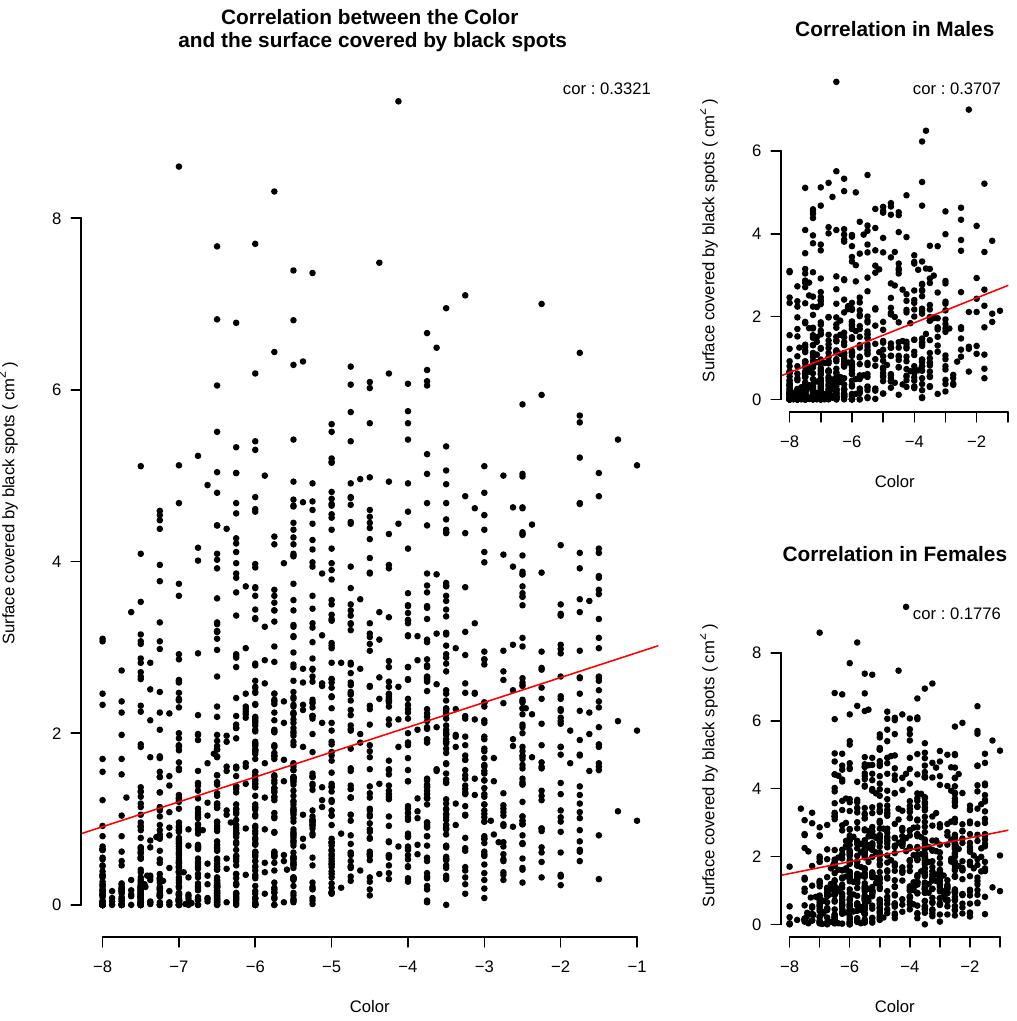

**Figure S2** – Correlation between the color of all the individuals and their spottiness (left panel), as well as within each sex (upper right for Males and bottom right for Females).

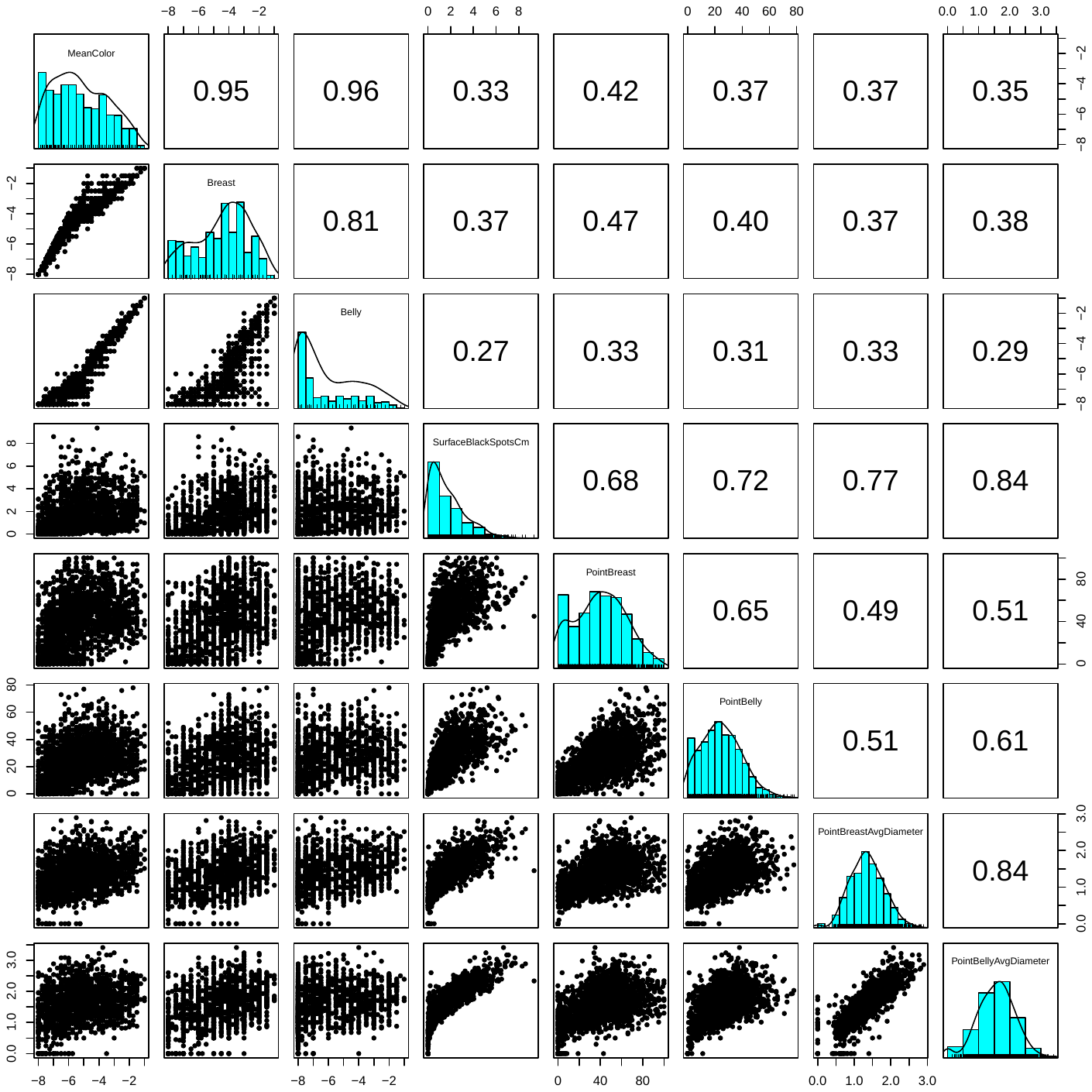

**Figure S3** – Distribution and correlation between all the color and spottiness variables. The three first rows and columns present the color variables (namely the “*MeanColor*” used in the main text, the “*Breast*” referring to the color of the breast of the individual and the “*belly*”, referring to the color of the belly of the individuals). The next five columns and rows resent the spottiness variables (namely “*SurfaceBackSpotsCm*” used as an overall measure of the spottiness in the main text, the “*PointBreast*” and “*PointBelly*” variables describing the number of Spots counted in each part of the body, and “*PointBreastAvgDiameter*” and “*PointBellyAvgDiameter*” presenting the average diameter of the spots in each body part).

#### Evolution of two melanin-based traits

**Table S1** – Choice of the best model explaining the evolution of the color (see Supplementary tables).

**Table S2** – Summary of the best model explaining the evolution of the color.

|  | **(MeanColor)** | | |
| --- | --- | --- | --- |
| *Predictors* | *Estimates* | *CI* | *p* |
| (Intercept) | -88.35 | -112.37 – -64.32 | **<0.001** |
| HatchYear | 0.04 | 0.03 – 0.05 | **<0.001** |
| Stage [Fledgling] | 0.51 | 0.43 – 0.59 | **<0.001** |
| Sex [2] | 1.37 | 1.30 – 1.44 | **<0.001** |
| **Random Effects** | | | |
| σ^2^ | 2.29 | | |
| τ_00_ _YearFirstObs_ | 0.07 | | |
| ICC | 0.03 | | |
| N _YearFirstObs_ | 30 | | |
| Observations | 7748 | | |
| Marginal R^2^ / Conditional R^2^ | 0.211 / 0.235 | | |

**Table S3** – Choice of the best model explaining the evolution of the spottiness (see Supplementary tables)

**Table S4** – Summary of the best model explaining the evolution of the spottiness.

|  | **(SurfaceBlackSpotsCm)** | | |
| --- | --- | --- | --- |
| *Predictors* | *Estimates* | *CI* | *p* |
| (Intercept) | -97.76 | -121.57 – -73.94 | **<0.001** |
| HatchYear | 0.05 | 0.04 – 0.06 | **<0.001** |
| Stage [Fledgling] | 0.27 | 0.17 – 0.36 | **<0.001** |
| Sex [2] | 0.91 | 0.83 – 0.99 | **<0.001** |
| **Random Effects** | | | |
| σ^2^ | 2.59 | | |
| τ_00_ _YearFirstObs_ | 0.07 | | |
| ICC | 0.03 | | |
| N _YearFirstObs_ | 30 | | |
| Observations | 6577 | | |
| Marginal R^2^ / Conditional R^2^ | 0.133 / 0.155 | | |

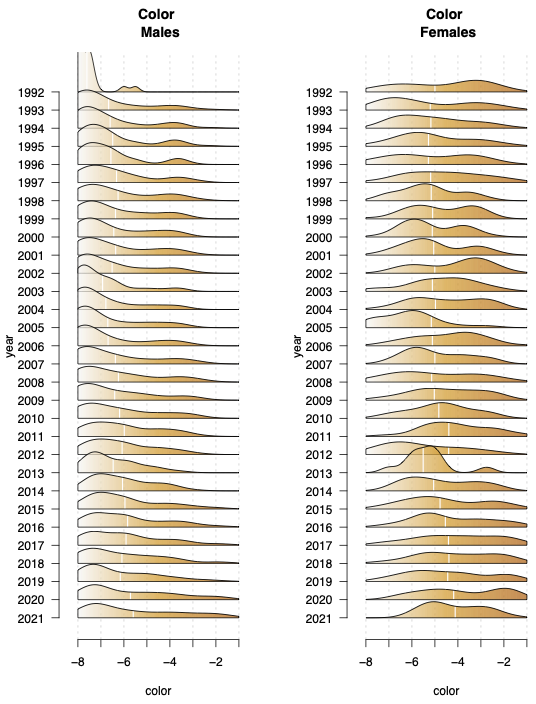

**Figure S4** - Evolution of the color of the individuals along the year for males (left) and females (right) based on adults alive that year. For each year, the white bar presents the mean of the distribution.

**Table S5** – Summary of the model explaining the evolution of the color in males only.

|  | **(MeanColor)** | | |
| --- | --- | --- | --- |
| *Predictors* | *Estimates* | *CI* | *p* |
| (Intercept) | -81.67 | -107.70 – -55.65 | **<0.001** |
| HatchYear | 0.04 | 0.02 – 0.05 | **<0.001** |
| Stage [Fledgling] | 0.59 | 0.47 – 0.72 | **<0.001** |
| **Random Effects** | | | |
| σ^2^ | 2.63 | | |
| τ_00_ _YearFirstObs_ | 0.07 | | |
| ICC | 0.03 | | |
| N _YearFirstObs_ | 30 | | |
| Observations | 3698 | | |
| Marginal R^2^ / Conditional R^2^ | 0.063 / 0.087 | | |

**Table S6** – Summary of the model explaining the evolution of the color in females only.

|  | **(MeanColor)** | | |
| --- | --- | --- | --- |
| *Predictors* | *Estimates* | *CI* | *p* |
| (Intercept) | -78.24 | -99.56 – -56.93 | **<0.001** |
| HatchYear | 0.04 | 0.03 – 0.05 | **<0.001** |
| Stage [Fledgling] | 0.45 | 0.35 – 0.55 | **<0.001** |
| **Random Effects** | | | |
| σ^2^ | 1.97 | | |
| τ_00_ _YearFirstObs_ | 0.06 | | |
| ICC | 0.03 | | |
| N _YearFirstObs_ | 30 | | |
| Observations | 4042 | | |
| Marginal R^2^ / Conditional R^2^ | 0.070 / 0.097 | | |

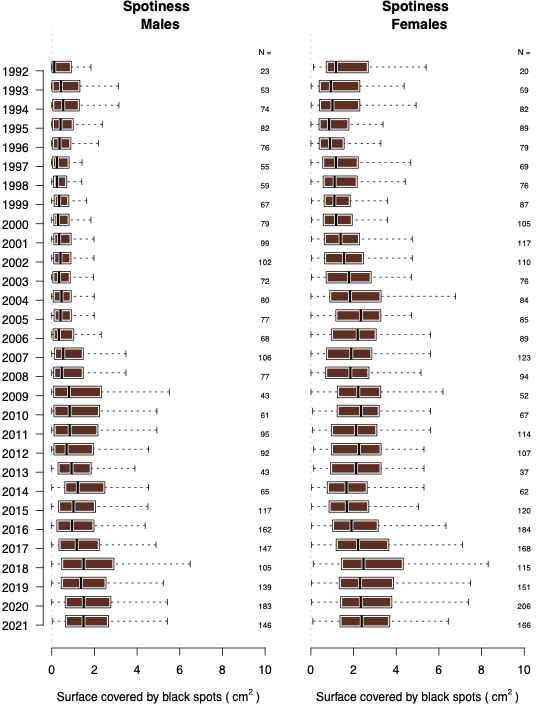

**Figure S5** - Evolution of the spottiness of the individuals along the year for males (left) and females (right) based on adults alive that year.

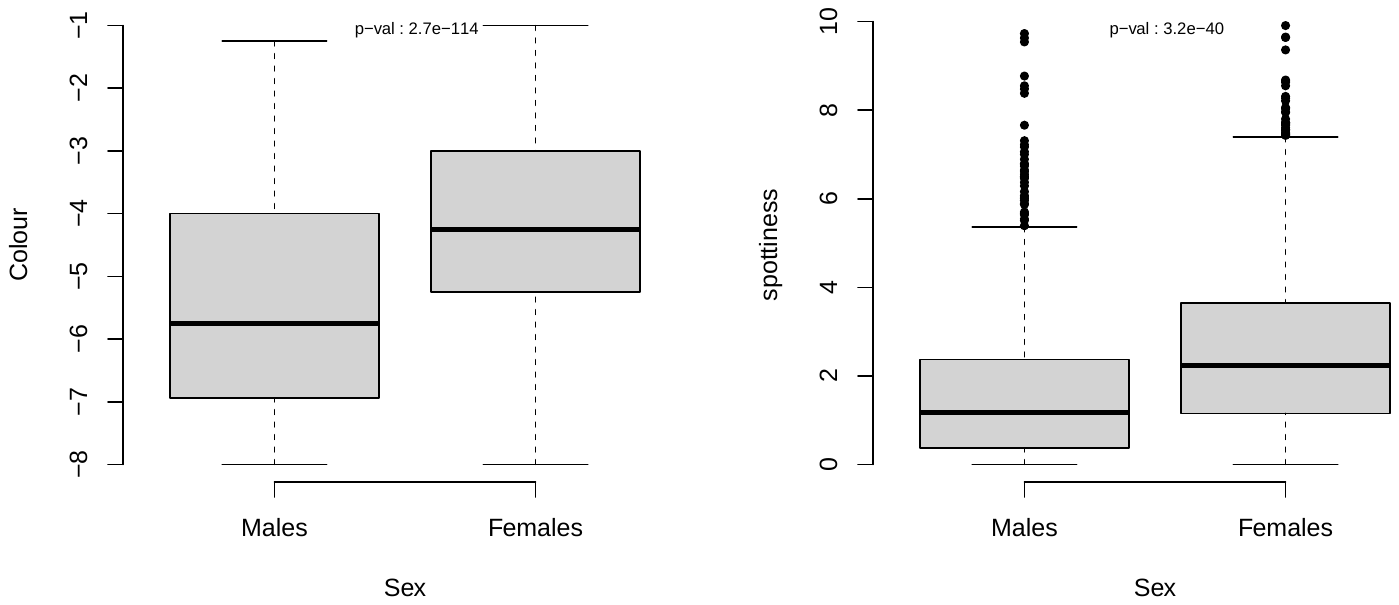

**Figure S6** – Sexual dimorphism in the color (left panel) and number of spots (right panel) in the barn owl. P-values on top of each plot present the result of the Wilcoxon Rank Sum test comparing both distributions.

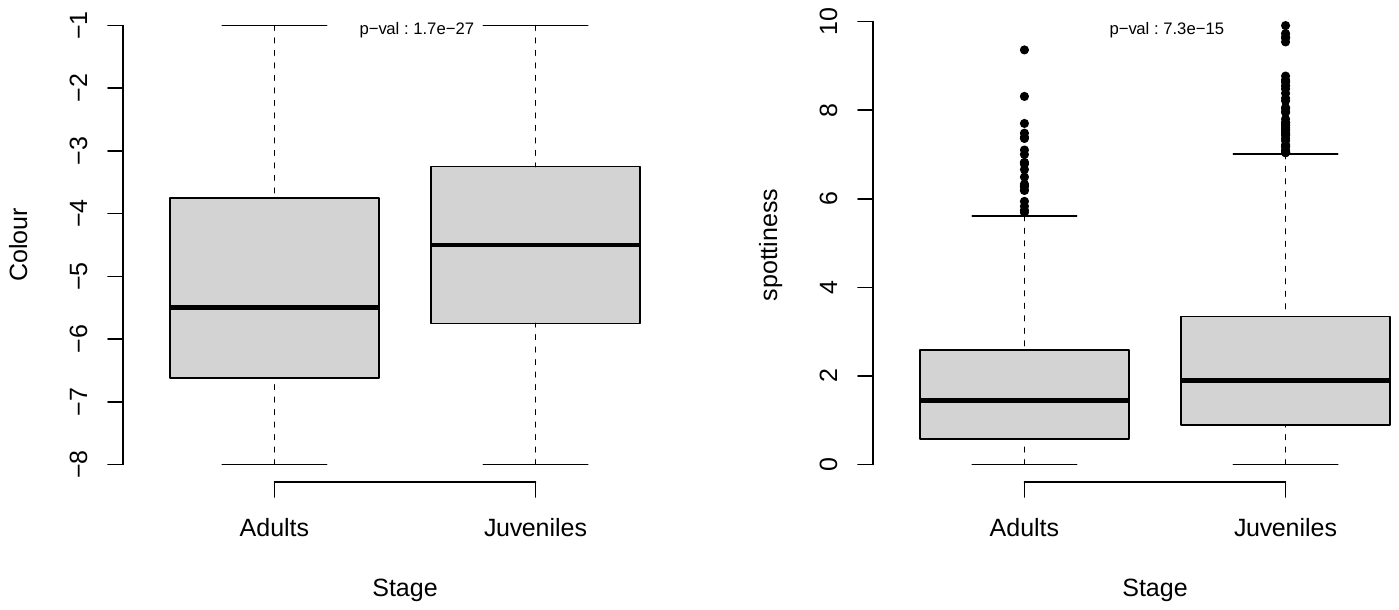

**Figure S7** – Difference between juveniles and adults in the color (left panel) and spottiness (right panel) in the barn owl. P-values on top of each plot present the result of the Wilcoxon Rank Sum test comparing both distributions.

**Table S7** – Summary of the model explaining the evolution of the spottiness in males only.

|  | **(SurfaceBlackSpotsCm)** | | |
| --- | --- | --- | --- |
| *Predictors* | *Estimates* | *CI* | *p* |
| (Intercept) | -109.56 | -135.66 – -83.47 | **<0.001** |
| HatchYear | 0.06 | 0.04 – 0.07 | **<0.001** |
| Stage [Fledgling] | 0.23 | 0.10 – 0.35 | **<0.001** |
| **Random Effects** | | | |
| σ^2^ | 2.07 | | |
| τ_00_ _YearFirstObs_ | 0.07 | | |
| ICC | 0.03 | | |
| N _YearFirstObs_ | 30 | | |
| Observations | 3118 | | |
| Marginal R^2^ / Conditional R^2^ | 0.105 / 0.136 | | |

**Table S8** – Summary of the model explaining the evolution of the spottiness in females only.

|  | **(SurfaceBlackSpotsCm)** | | |
| --- | --- | --- | --- |
| *Predictors* | *Estimates* | *CI* | *p* |
| (Intercept) | -97.76 | -121.57 – -73.94 | **<0.001** |
| HatchYear | 0.05 | 0.04 – 0.06 | **<0.001** |
| Stage [Fledgling] | 0.27 | 0.17 – 0.36 | **<0.001** |
| Sex [2] | 0.91 | 0.83 – 0.99 | **<0.001** |
| **Random Effects** | | | |
| σ^2^ | 2.59 | | |
| τ_00_ _YearFirstObs_ | 0.07 | | |
| ICC | 0.03 | | |
| N _YearFirstObs_ | 30 | | |
| Observations | 6577 | | |
| Marginal R^2^ / Conditional R^2^ | 0.133 / 0.155 | | |

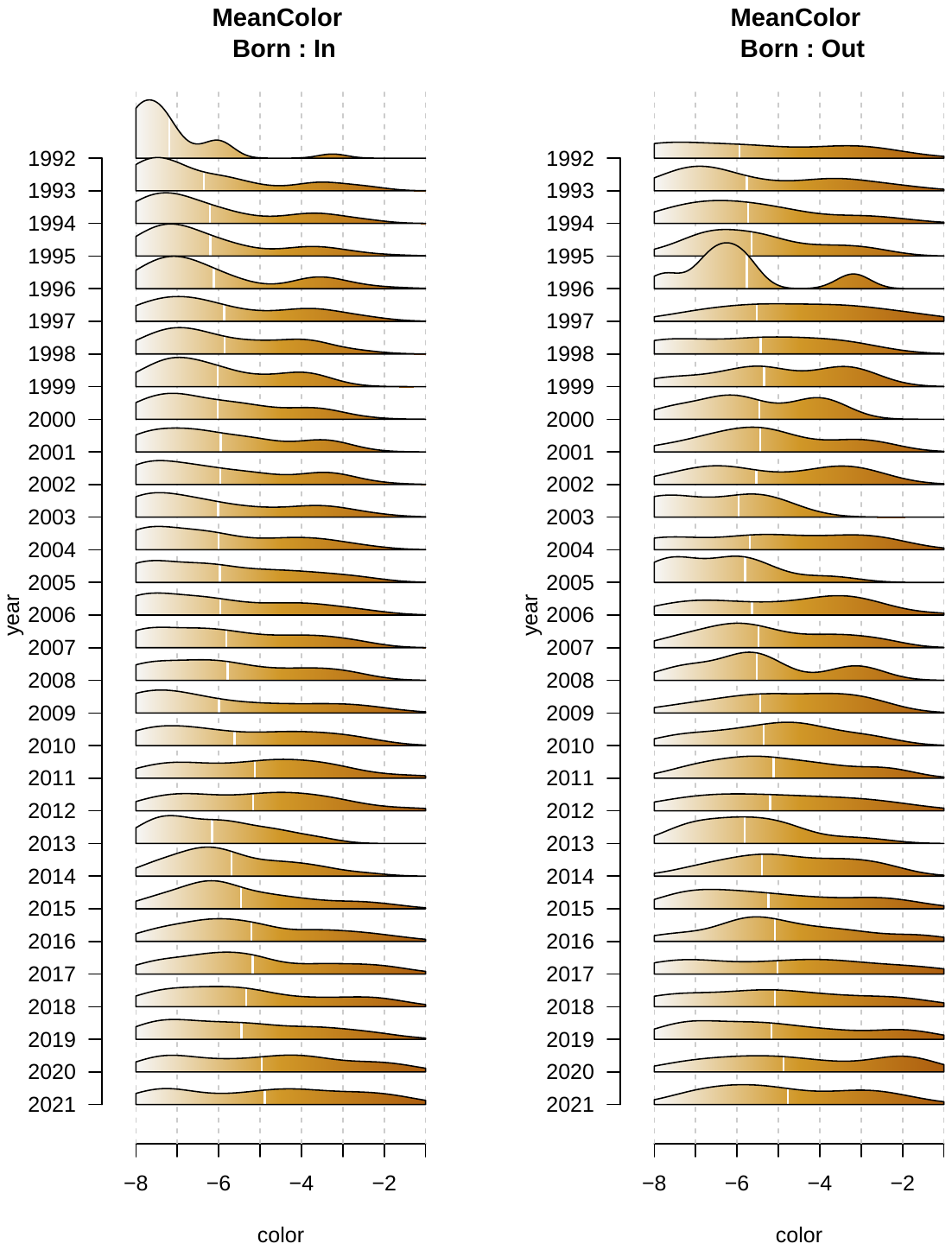

**Figure S8** – Evolution of the color of the individuals along the year for individuals born in the system (i.e. ringed as fledglings - left panel) or outside the system (i.e. ringed as adults - left panel). Distributions are based on individuals alive that year.

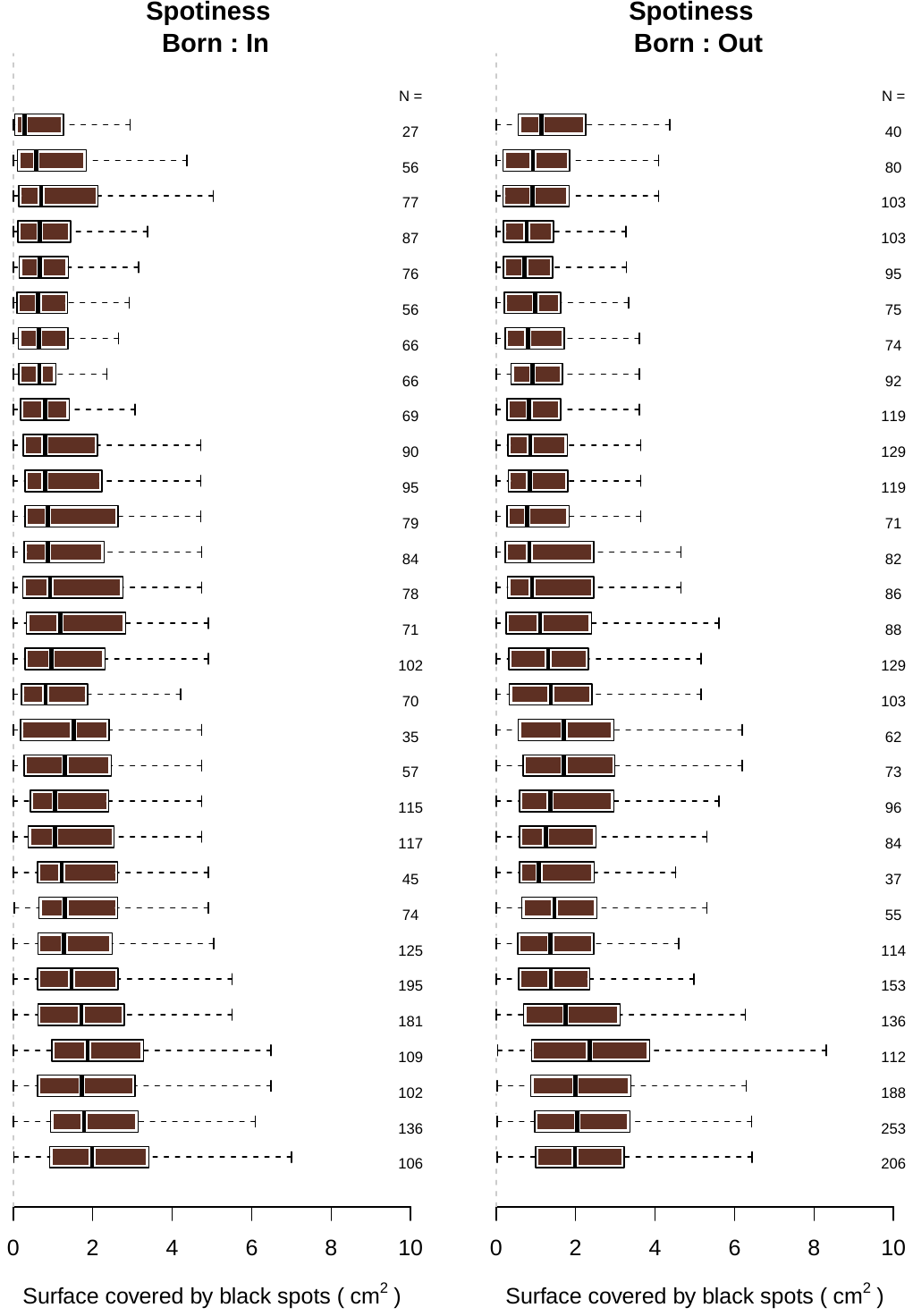

**Figure S9** - Evolution of the Spottiness of the individuals along the year for individuals born in the system (i.e. ringed as fledgelings - left panel) or outside the system (i.e. ringed as adults - left panel). Distributions are based on individuals alive that year.

### Population-wide whole genome sequencing of the Swiss population

#### Genetic diversity in reference panel

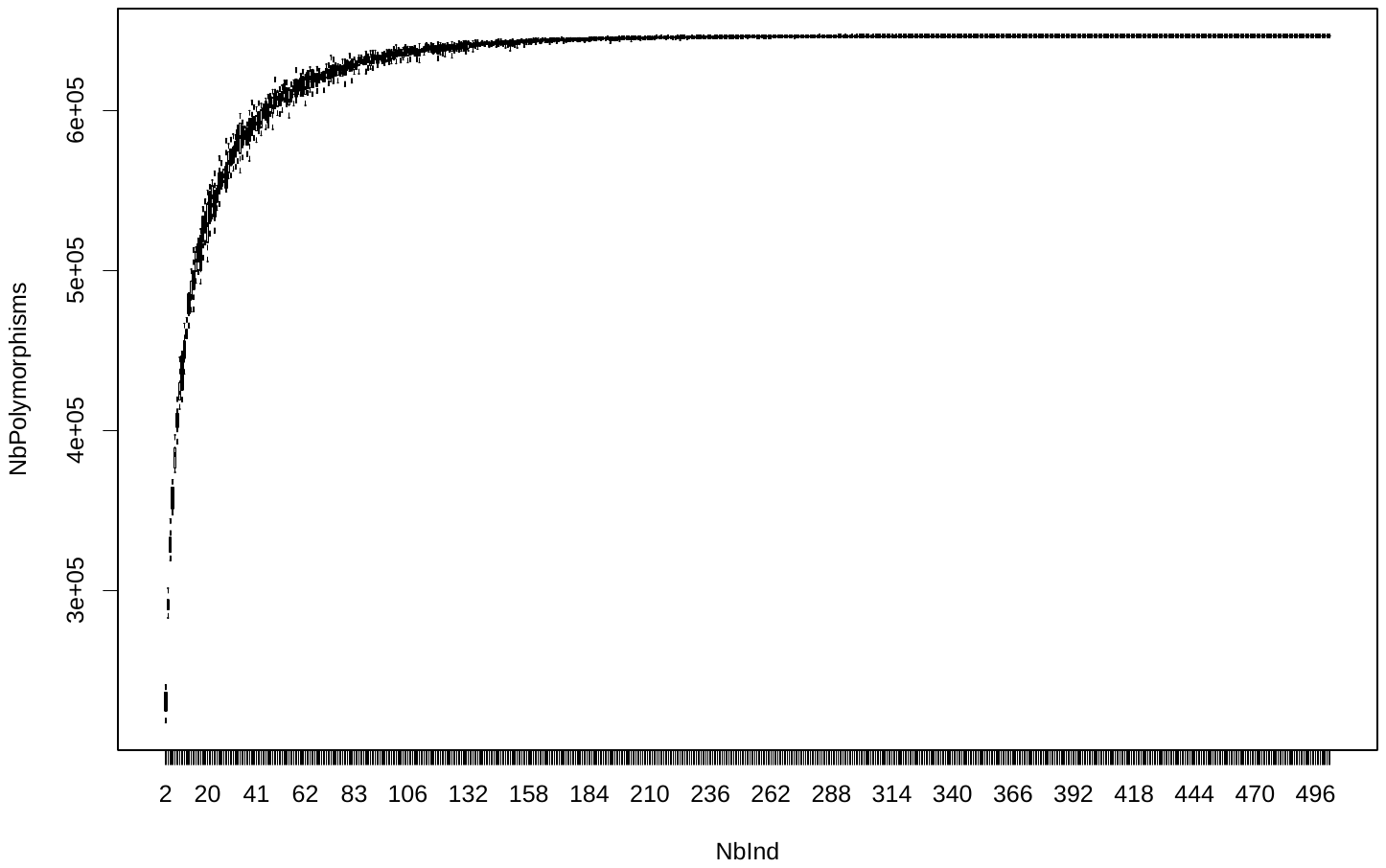

**Figure S10** – Relation between the number of individuals composing the reference panel and the number of variants identified. Each boxplot presents the results for ten random draws of *x* individuals, for which we recorded the number of polymorphic variant (each allele was present at least one time). With a sample size of ~180 and more individuals, the number of variants does not increase anymore, thus implying that the reference panel capture well the diversity of frequent alleles (see M&M section for filtering details).

#### Phasing of the reference panel

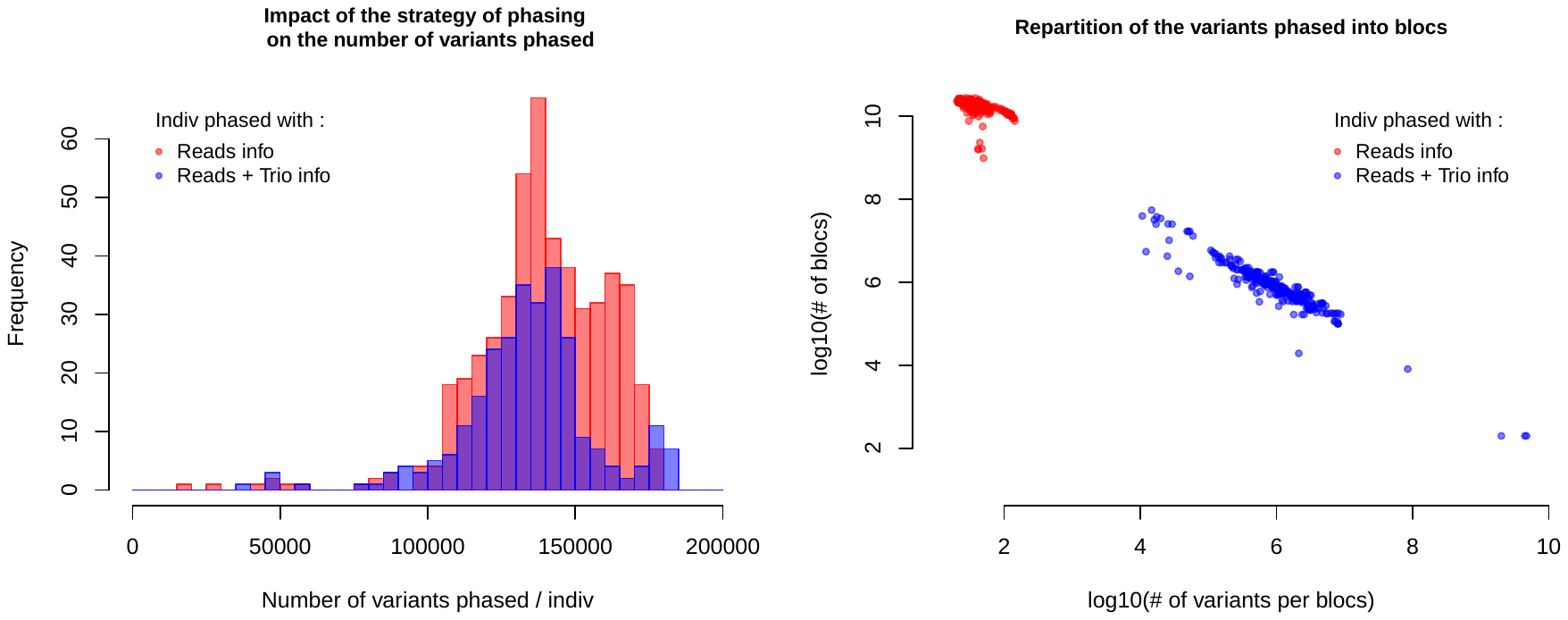

**Figure S11** – results of the read-based phasing (using WhatsHap) of the reference panel. The left panel depict the number of variants phased per individuals, with in red individuals phased only with their read information and in blue the individuals with a phasing done using both reads and trio information (see M&M section for details). Both strategies perform equally, phasing a high number of variants per individuals. The right panel present the relation between the number of blocs of variants phased in the same individuals and the size of the number of variants par blocs. The Color code is the same as before, based on the phasing data available. Individuals phased only with the reads have a higher number of blocs with less variants per blocs relative to the individuals phased taking advantage of the trio information on top of the reads.

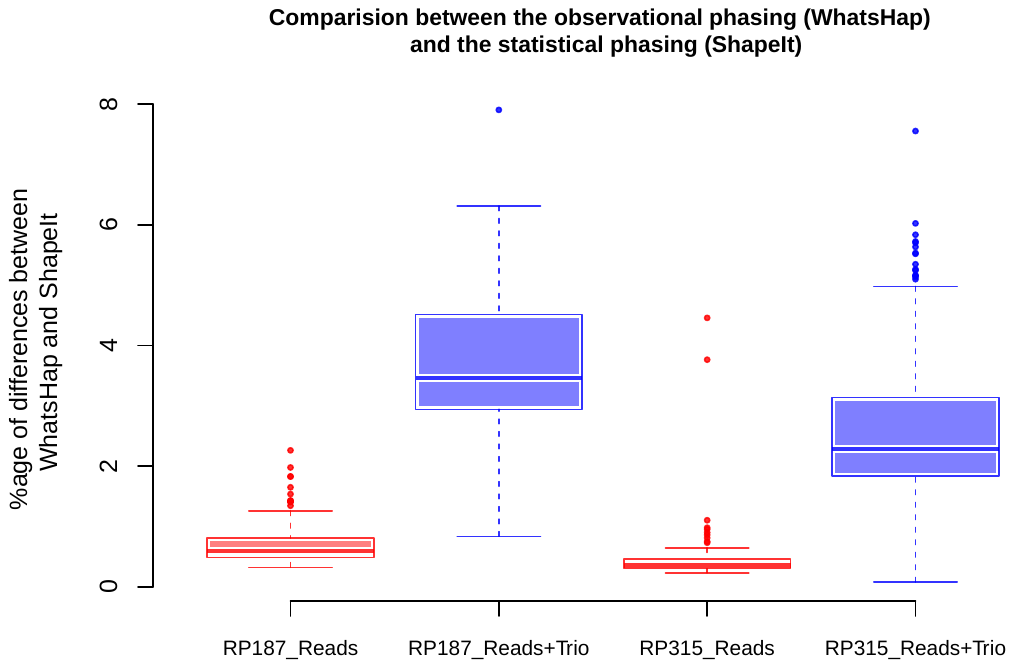

**Figure S12** – Boxplots presenting the percentage of differences between the phasing based on the reads (and trio) information and the statistical phasing. Each value of the boxplot represents the differences between two independent phasing of the same individual, one with the read based approach only and with the statistical approach only. Individuals phased with the trio information have longer blocs with more variants, thus increasing the proportion of blocs with errors compare to individuals phased only with reads.

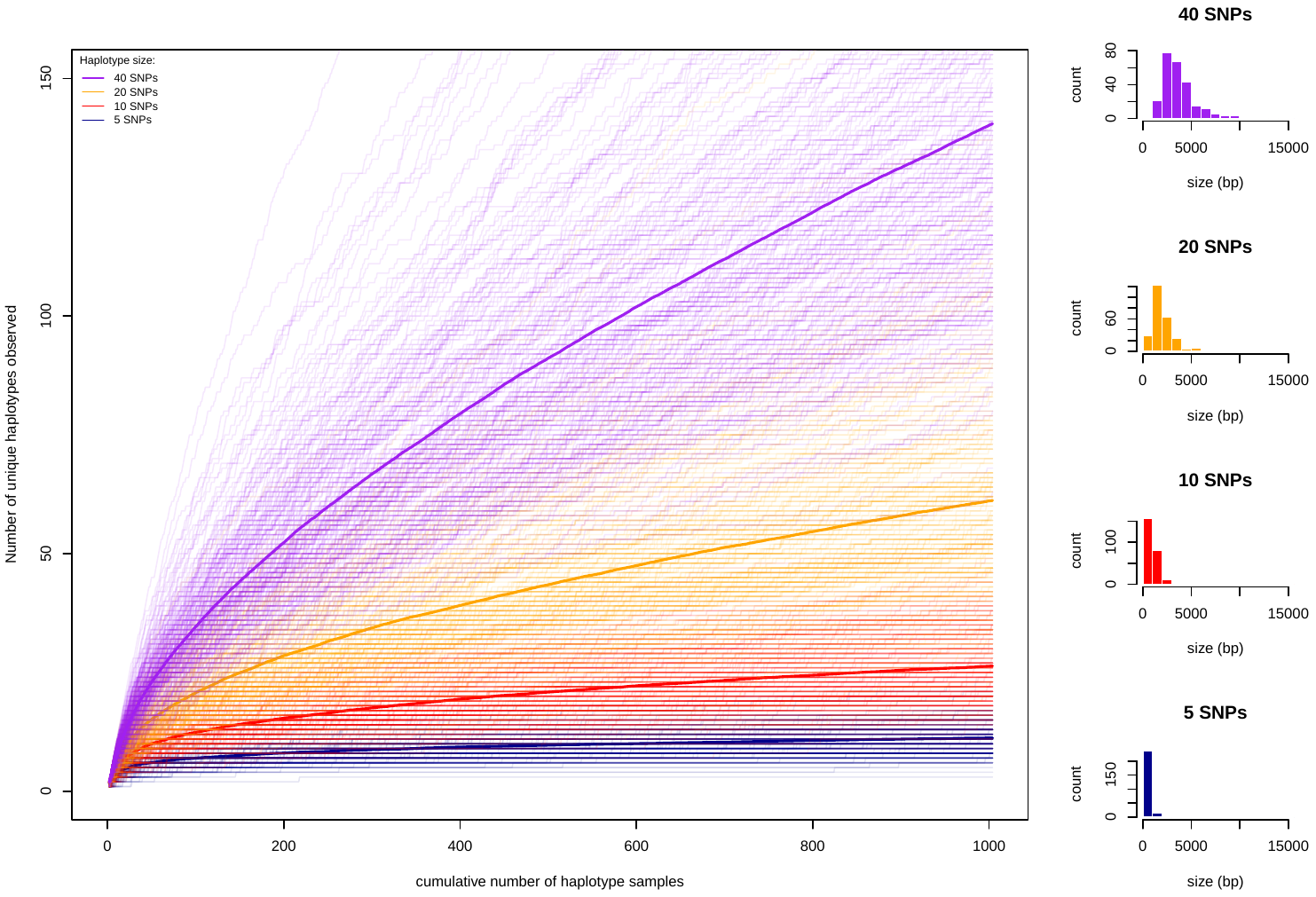

**Figure S13** - Relation between the number of local haplotypes composing the reference panel and the number of unique haplotypes for four different haplotype size (5 SNPs, 10 SNPs, 20 SNPs and 40 SNPs). Each line represents the cumulative number of unique haplotypes within one random subsampling of all haplotypes present in each region. Regions were selected randomly along the genome. Solid lines represent the mean cumulative number of haplotypes over 200 random regions. The right panel present the distribution of the sizes of the regions randomly selected and presented in the left panel.

#### Low coverage sequencing of 2778 owls

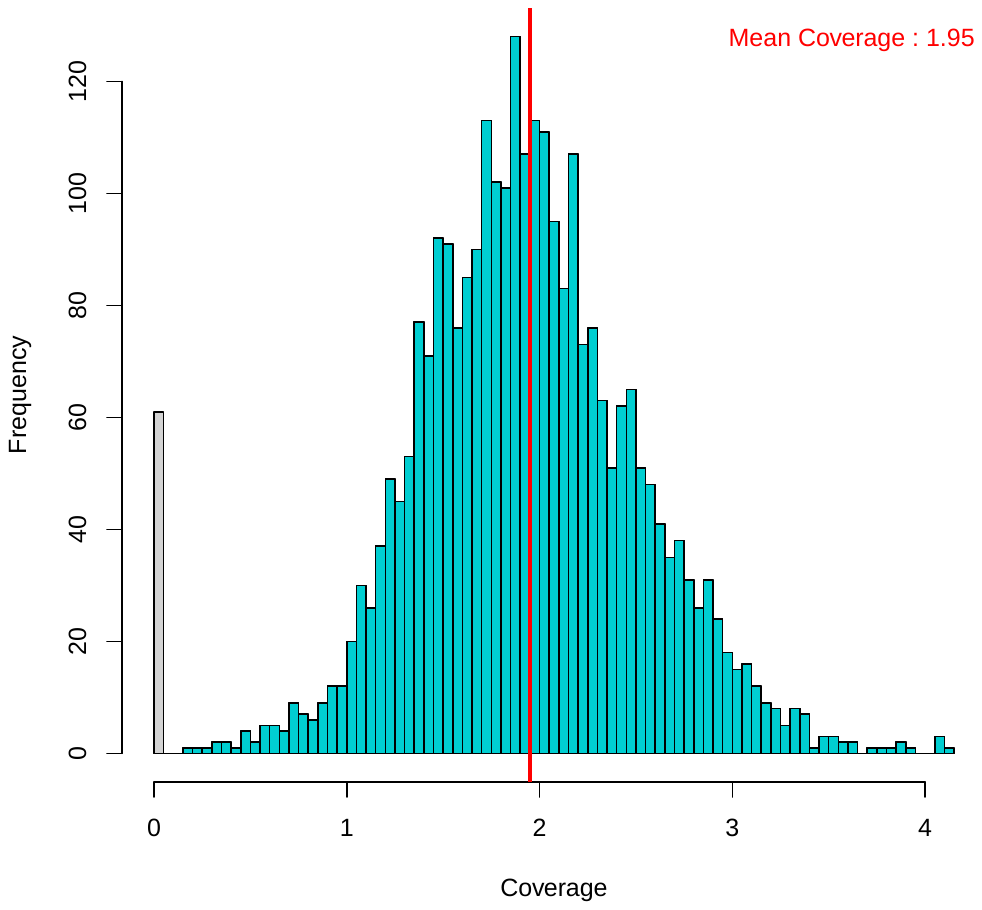

**Figure S14** – Depth of coverage of the 2778 owls sequenced at low coverage. The grey at 0 represent the coverage of the control wells used during sequencing (see M&M section for details). Turquoise distribution represents the coverage of the owls. Red vertical line depicts the mean value of the distribution at 1.95X. Minimum coverage: 0.2X. Maximum Coverage: 4.15X.

### Heritability of the coloration and the Spottiness

**Table S9** - Summary of the model estimating the heritability of the coloration.

Family: gaussian

Links: mu = identity; sigma = identity

Formula: MeanColor ~ 1 + as.factor(StageBin) + as.factor(SexWGS) + (1 | gr(ID, cov = grm))

Data: TabSummary (Number of observations: 3083)

Draws: 4 chains, each with iter = 5000; warmup = 2000; thin = 1; total post-warmup draws = 12000

Group-Level Effects: ~ID (Number of levels: 3083)

Estimate	Est.Error	l-95% CI	u-95% CI	Rhat	Bulk_ESS	Tail_ESS

sd(Intercept)	1.81	0.06	1.70	1.93	1.01	784	1743

Population-Level Effects:

Estimate	Est.Error	l-95% CI	u-95% CI	Rhat	Bulk_ESS	Tail_ESS

Intercept	-5.79	0.10	-5.98	-5.60	1.00	5037	6394

as.factorStageBin2	0.73	0.04	0.64	0.81	1.00	9996	9728

as.factorSexWGS2	1.24	0.12	1.00	1.48	1.00	4793	6197

Family Specific Parameters:

Estimate	Est.Error	l-95% CI	u-95% CI	Rhat	Bulk_ESS	Tail_ESS

sigma	0.86	0.03	0.79	0.92	1.00	709	1389

**Table S10** – Summary of the model estimating the heritability of the spottiness.

Family: gaussian

Links: mu = identity; sigma = identity

Formula: SurfaceBlackSpotsCm ~ 1 + as.factor(Stage) + as.factor(Sex) + (1 | gr(ID, cov = grm))

Draws: 4 chains, each with iter = 5000; warmup = 2000; thin = 1; total post-warmup draws = 12000

Group-Level Effects:

~ID (Number of levels: 2480)

Estimate	Est.Error	l-95% CI	u-95% CI	Rhat	Bulk_ESS	Tail_ESS

sd(Intercept)	1.97	0.07	1.82	2.10	1.01	848	2175

Population-Level Effects:

Estimate	Est.Error	l-95% CI	u-95% CI	Rhat	Bulk_ESS	Tail_ESS

Intercept	1.24	0.11	1.02	1.46	1.00	3612	5603

as.factorStageBin2	0.47	0.06	0.36	0.58	1.00	7582	8424

as.factorSexWGS2	1.03	0.14	0.76	1.29	1.00	3678	5746

Family Specific Parameters:

Estimate	Est.Error	l-95% CI	u-95% CI	Rhat	Bulk_ESS	Tail_ESS

sigma	0.99	0.04	0.91	1.06	1.01	759	1937

### Genome Wide Association Study

#### GWAS – Plumage coloration

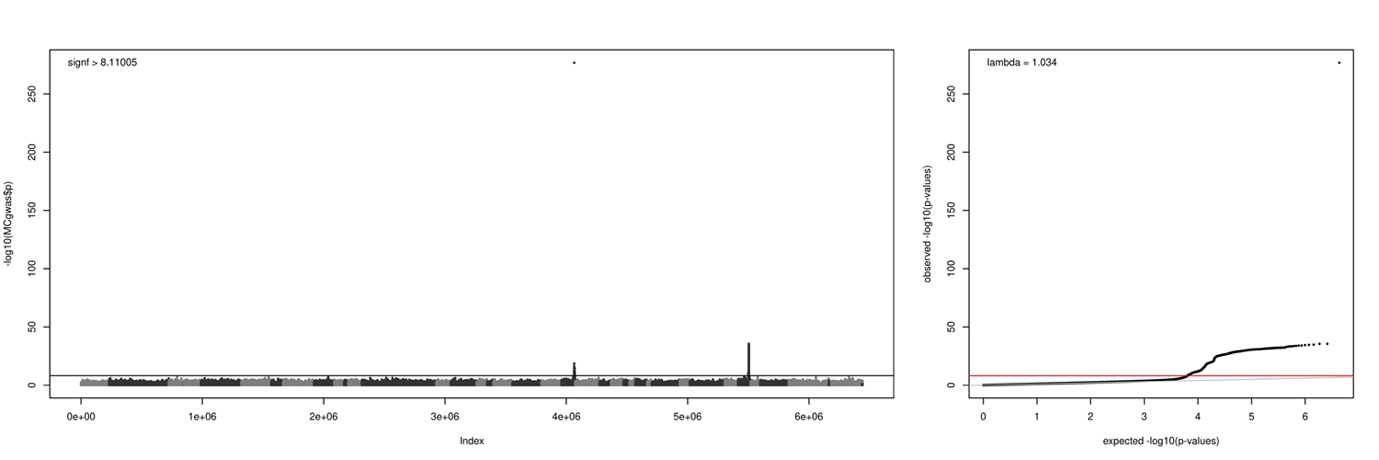

**Figure S15** – Genome Wide Association (GWA) study between the genotypes and the **plumage coloration of the entire ventral part**. Left panel: Association scores (-log_10_[p-values]) of each SNPs with the trait. Alternated colors depict the successive super-scaffolds (SC_) in the genome. Horizontal line presents the significance threshold according to Bonferroni. Right panel: QQ-plot of the GWAS presented on the left panel. Solid grey line represents the 1:1 line.

**Table S11 –** List of SNPs significant in the Genome Wide Association (GWA) study between the genotypes and the **plumage coloration of the entire ventral part.**

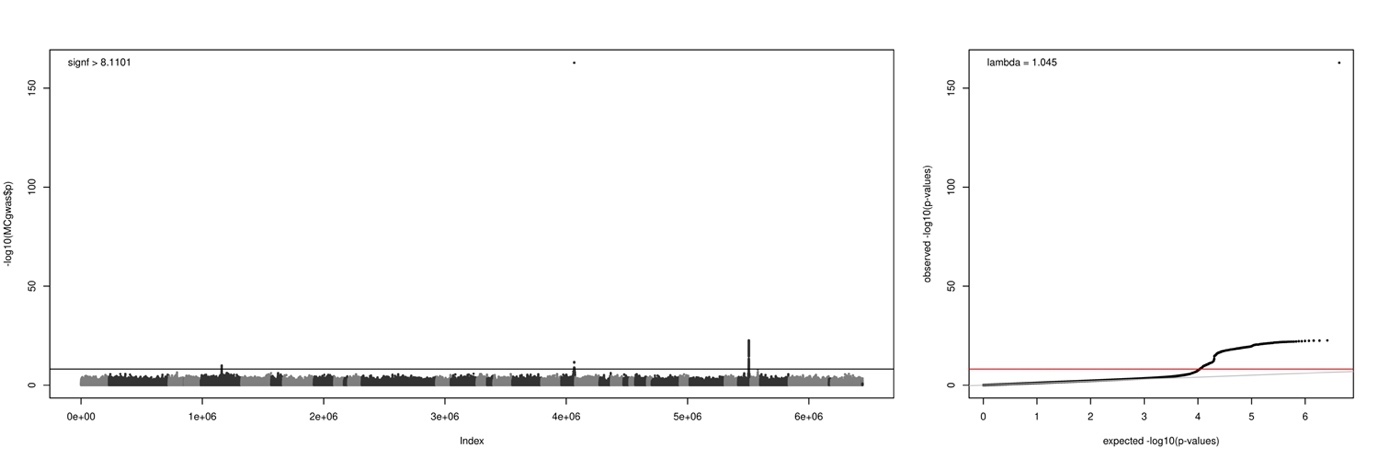

**Figure S16** – Genome Wide Association (GWA) study between the genotypes and the **plumage coloration of the breast**. Left panel: Association scores (-log_10_[p-values]) of each SNPs with the trait. Alternated colors depict the successive super-scaffolds (SC_) in the genome. Horizontal line presents the significance threshold according to Bonferroni. Right panel: QQ-plot of the GWAS presented on the left panel. Solid grey line represents the 1:1 line.

**Table S12 –** List of SNPs significant in the Genome Wide Association (GWA) study between the genotypes and the **plumage coloration of the breast.**

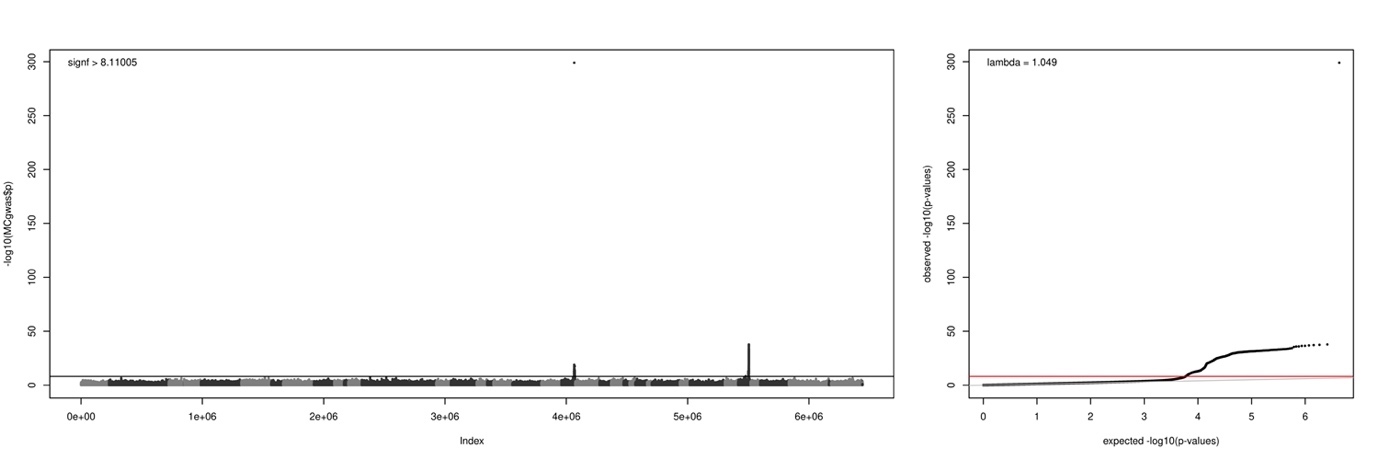

**Figure S17** – Genome Wide Association (GWA) study between the genotypes and the **plumage coloration of the belly**. Left panel: Association scores (-log_10_[p-values]) of each SNPs with the trait. Alternated colors depict the successive super-scaffolds (SC_) in the genome. Horizontal line presents the significance threshold according to Bonferroni. Right panel: QQ-plot of the GWAS presented on the left panel. Solid grey line represents the 1:1 line.

**Table S13 –** List of SNPs significant in the Genome Wide Association (GWA) study between the genotypes and the **plumage coloration of the belly.**

#### GWAS- ventral spottiness

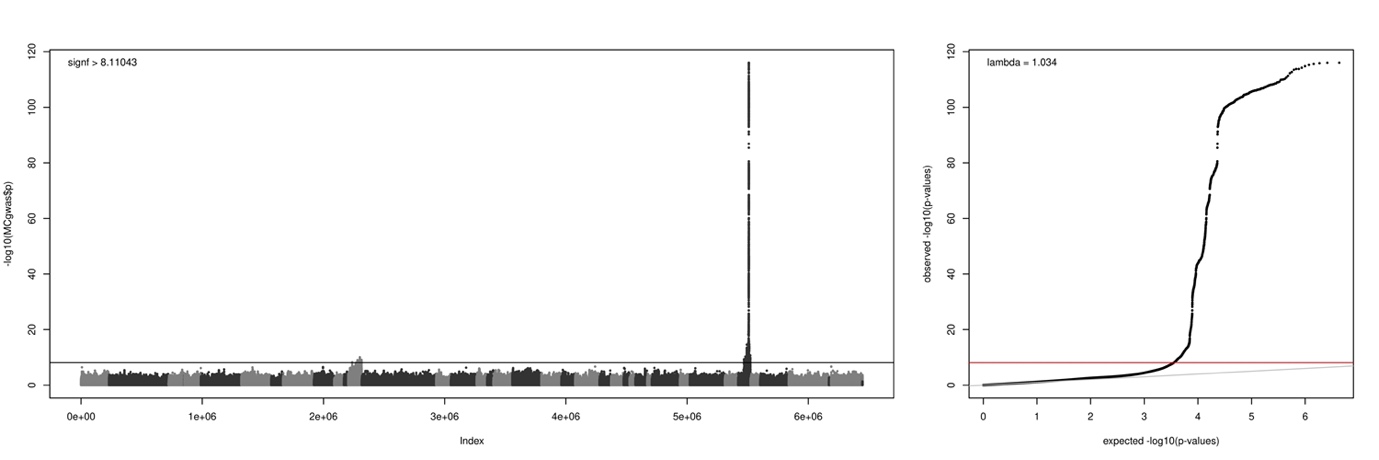

**Figure S18** – Genome Wide Association (GWA) study between the genotypes and the **ventral spottiness**. Left panel: Association scores (-log_10_[p-values]) of each SNPs with the trait. Alternated colors depict the successive super-scaffolds (SC_) in the genome. Horizontal line presents the significance threshold according to Bonferroni. Right panel: QQ-plot of the GWAS presented on the left panel. Solid grey line represents the 1:1 line.

**Table S14 –** List of SNPs significant in the Genome Wide Association (GWA) study between the genotypes and the **ventral spottiness.**

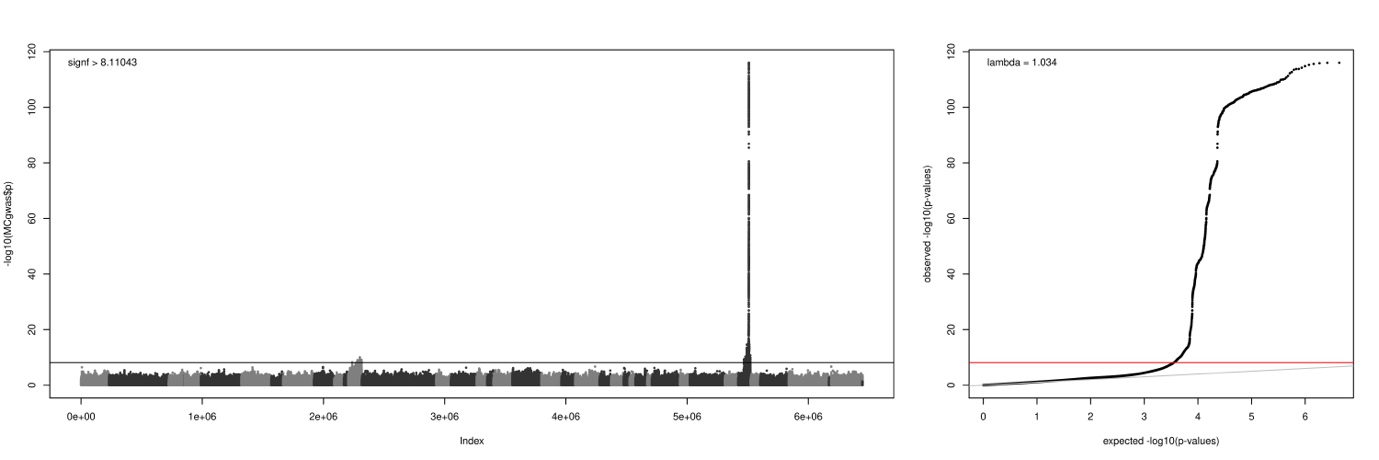

**Figure S19** – Genome Wide Association (GWA) study between the genotypes and the mean **diameter of the spots**. Left panel: Association scores (-log_10_[p-values]) of each SNPs with the trait. Alternated colors depict the successive super-scaffolds (SC_) in the genome. Horizontal line presents the significance threshold according to Bonferroni. Right panel: QQ-plot of the GWAS presented on the left panel. Solid grey line represents the 1:1 line.

**Table S15 –** List of SNPs significant in the Genome Wide Association (GWA) study between the genotypes and the mean **diameter of the spots.**

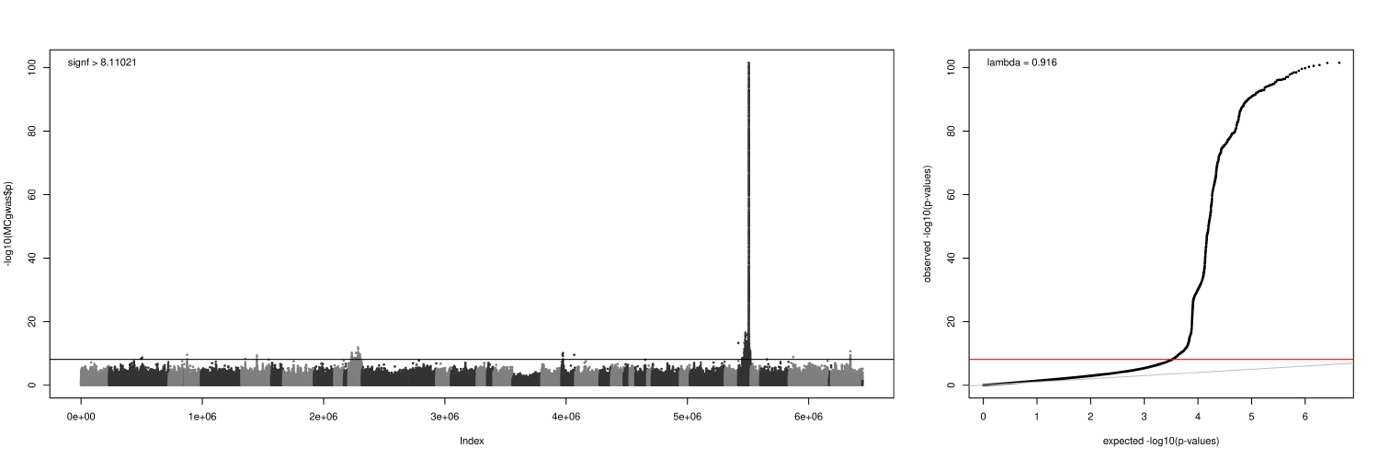

**Figure S20** – Genome Wide Association (GWA) study between the genotypes and the **number of spots**. Left panel: Association scores (-log_10_[p-values]) of each SNPs with the trait. Alternated colors depict the successive super-scaffolds (SC_) in the genome. Horizontal line presents the significance threshold according to Bonferroni. Right panel: QQ-plot of the GWAS presented on the left panel. Solid grey line represents the 1:1 line.

**Table S16 –** List of SNPs significant in the Genome Wide Association (GWA) study between the genotypes and the **number of spots.**

### Differential gene expression links genotype to phenotype

#### Gene expression and RT-qPCR assay

**Table S17 -** Sequences and efficiency of the probe and primer pairs used to quantify the gene present on the Z chromosome.

#### Genotype, gene expression and phenotype

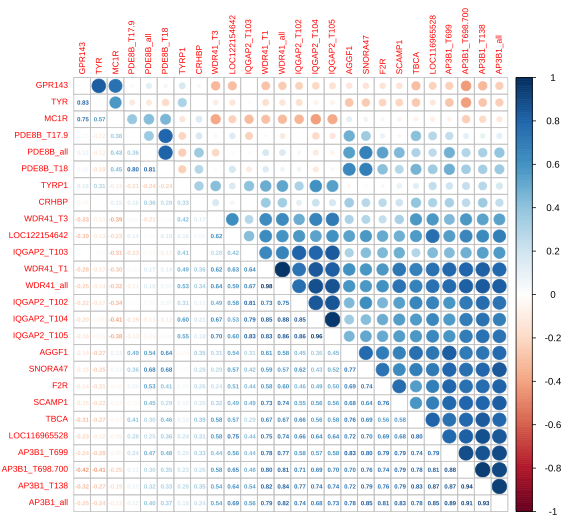

**Figure S21** - Pearson’s correlation between the relative normalised sqrt-root expression of all the transcripts of the tested genes.

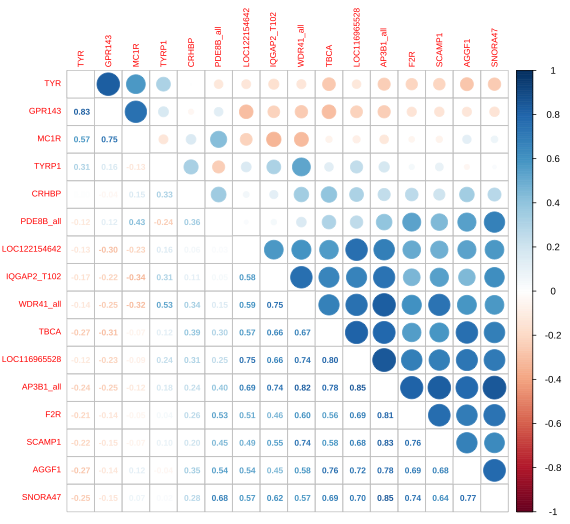

**Figure S22** - Pearson’s correlation between the relative gene expression for the primers-probe pairs represented either all transcripts of a gene or the most expressed transcript for IQGAP2.

**Table S18** - Table comparing models explaining the expression without and with the genotype ((formula: "sqrt(Gene) ~ 1, random = ~1 | nestbox") and (formula: "sqrt(Gene) ~ as.factor(GenoZ), random = ~1 | nestbox")) for males (left) and females (right). Significant values after Bonferroni correction are highlighted in bold. BF stands for Bonferroni-corrected.

| Gene | p-value_Males | p-value_Males_BF | p-value_Females | p-value_Females_BF |
| --- | --- | --- | --- | --- |
| AGGF1 | 0.6059 | 1 | 0.1919 | 1 |
| AP3B1_all | 0.1874 | 1 | 0.2386 | 1 |
| CRHBP | 0.4339 | 1 | 0.5641 | 1 |
| F2R | 0.8578 | 1 | 0.1296 | 1 |
| IQGAP2_T102 | 0.2778 | 1 | 0.251 | 1 |
| LOC116965528 | 0.0882 | 1 | 0.4919 | 1 |
| LOC122154642 | 0.0424 | 0.5088 | 0.2591 | 1 |
| PDE8B_all | **0.0018** | **0.0216** | 0.7921 | 1 |
| SCAMP1 | 0.2601 | 1 | 0.6906 | 1 |
| SNORA47 | 0.2833 | 1 | 0.2973 | 1 |
| TBCA | 0.2472 | 1 | 0.8725 | 1 |
| WDR41_all | 0.0783 | 0.9396 | **0.0025** | **0.03** |

**Table S19** – Summary of the model linking the genotype and PDE8B expression in males.

|  | **sqrt(PDE8B_all)** | | |
| --- | --- | --- | --- |
| *Predictors* | *Estimates* | *CI* | *p* |
| (Intercept) | 1.06 | 0.92 – 1.20 | **<0.001** |
| GenoZ [CT] | -0.34 | -0.59 – -0.08 | **0.032** |
| GenoZ [TT] | -0.28 | -0.46 – -0.10 | **0.009** |
| **Random Effects** | | | |
| σ^2^ | 0.02 | | |
| τ_00_ _nestbox_ | 0.00 | | |
| N _nestbox_ | 15 | | |
| Observations | 19 | | |
| Marginal R^2^ / Conditional R^2^ | 0.499 / NA | | |

**Table S20** – Summary of the model linking the genotype and PDE8B expression in females.

|  | **sqrt(PDE8B_all)** | | |
| --- | --- | --- | --- |
| *Predictors* | *Estimates* | *CI* | *p* |
| (Intercept) | 0.56 | 0.39 – 0.73 | **<0.001** |
| GenoZ [T] | -0.02 | -0.23 – 0.18 | 0.813 |
| **Random Effects** | | | |
| σ^2^ | 0.02 | | |
| τ_00_ _nestbox_ | 0.00 | | |
| N _nestbox_ | 12 | | |
| Observations | 13 | | |
| Marginal R^2^ / Conditional R^2^ | 0.006 / NA | | |

**Table S21** – Summary of the model linking the genotype and WDR41 expression in males.

|  | **sqrt(WDR41_all)** | | |
| --- | --- | --- | --- |
| *Predictors* | *Estimates* | *CI* | *p* |
| (Intercept) | 0.93 | 0.79 – 1.07 | **<0.001** |
| GenoZ [CT] | 0.03 | -0.23 – 0.29 | 0.753 |
| GenoZ [TT] | -0.13 | -0.31 – 0.05 | 0.174 |
| **Random Effects** | | | |
| σ^2^ | 0.02 | | |
| τ_00_ _nestbox_ | 0.00 | | |
| N _nestbox_ | 15 | | |
| Observations | 19 | | |
| Marginal R^2^ / Conditional R^2^ | 0.245 / NA | | |

**Table S22** – Summary of the model linking the genotype and WDR41 expression in females.

|  | **sqrt(WDR41_all)** | | |
| --- | --- | --- | --- |
| *Predictors* | *Estimates* | *CI* | *p* |
| (Intercept) | 0.71 | 0.66 – 0.76 | **<0.001** |
| GenoZ [T] | -0.10 | -0.16 – -0.04 | **0.007** |
| **Random Effects** | | | |
| σ^2^ | 0.00 | | |
| τ_00_ _nestbox_ | 0.00 | | |
| N _nestbox_ | 12 | | |
| Observations | 13 | | |
| Marginal R^2^ / Conditional R^2^ | 0.525 / NA | | |

**Table S23** – Summary of the model for linking PDE8B expression to spottiness.

|  | **SurfaceBlackSpotsCm** | | |
| --- | --- | --- | --- |
| *Predictors* | *Estimates* | *CI* | *p* |
| (Intercept) | 178.84 | 79.53 – 278.15 | **0.002** |
| PDE8B all [sqrt] | -136.09 | -256.42 – -15.77 | **0.037** |
| Sex [2] | 51.91 | -0.87 – 104.70 | 0.064 |
| **Random Effects** | | | |
| σ^2^ | 1866.17 | | |
| τ_00_ _nestbox_ | 2991.91 | | |
| ICC | 0.62 | | |
| N _nestbox_ | 20 | | |
| Observations | 32 | | |
| Marginal R^2^ / Conditional R^2^ | 0.342 / 0.747 | | |

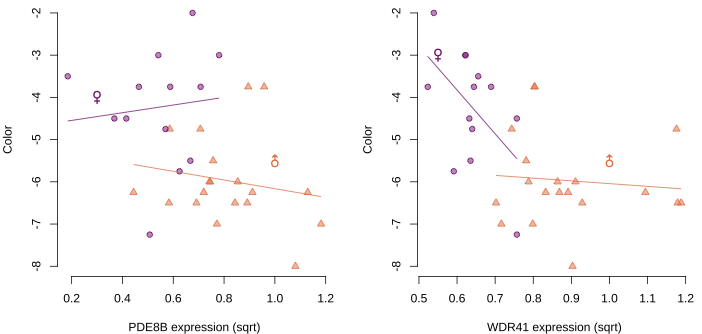

**Figure S22** – Relation between PDE8B expression (left) and WDR41 (right) expression and the color of the individuals. Lines represent the linear modelling of the relation within each sex.

### Sexual dimorphism and sex specific genetic architecture

#### Genetic architecture of coloration

**Table S24** **–** Selection of the best model explaining the color in males.

**Table S25** – Summary of the model best model for the genetic architecture of the coloration in males.

Family: gaussian

Links: mu = identity; sigma = identity

Formula: MeanColor ~ 1 + as.factor(StageBin) + as.factor(GenoMC1R) + as.factor(GenoZcol) + as.factor(GenoCorin) + (1 | gr(ID, cov = grm))

Data: TabSummary[TabSummary$SexWGS == 1, ] (Number of observations: 1480)

Draws: 4 chains, each with iter = 5000; warmup = 2000; thin = 1; total post-warmup draws = 12000

Group-Level Effects:

~ID (Number of levels: 1480)

Estimate	Est.Error	l-95% CI	u-95% CI	Rhat	Bulk_ESS	Tail_ESS

sd(Intercept)	1.03	0.07	0.90	1.16	1.00	1074	2364

Population-Level Effects:

Estimate	Est.Error	l-95% CI	u-95% CI	Rhat	Bulk_ESS	Tail_ESS

**Intercept	-7.34	0.11	-7.55	-7.13	1.00	6001	7850**

**as.factorStageBin2	0.67	0.05	0.57	0.77	1.00	9863	9330**

**as.factorGenoMC1R1	2.49	0.06	2.37	2.61	1.00	8210	8834**

**as.factorGenoMC1R2	3.40	0.15	3.10	3.69	1.00	8106	8927**

**as.factorGenoZcol1	0.34	0.07	0.21	0.47	1.00	6032	8199**

**as.factorGenoZcol2	0.93	0.09	0.76	1.11	1.00	5485	7036**

**as.factorGenoCorin1	0.28	0.07	0.13	0.42	1.00	6234	8319**

**as.factorGenoCorin2	0.37	0.09	0.19	0.54	1.00	6078	8086**

Family Specific Parameters:

Estimate	Est.Error	l-95% CI	u-95% CI	Rhat	Bulk_ESS	Tail_ESS

sigma	0.73	0.03	0.67	0.80	1.00	1103	2037

**Table S26** – Selection of the best model explaining the color in females.

**Table S27** – Summary of the model best model for the genetic architecture of the coloration in females.

Family: gaussian

Links: mu = identity; sigma = identity

Formula: MeanColor ~ 1 + as.factor(StageBin) + as.factor(GenoMC1R) + as.factor(GenoZcol) + as.factor(GenoCorin) + as.factor(GenoMC1R * GenoZcol) + (1 | gr(ID, cov = grm))

Data: TabSummary[TabSummary$SexWGS == 2, ] (Number of observations: 1603)

Draws: 4 chains, each with iter = 5000; warmup = 2000; thin = 1; total post-warmup draws = 12000

Group-Level Effects: ~ID (Number of levels: 1603)

Estimate	Est.Error	l-95% CI	u-95% CI	Rhat	Bulk_ESS	Tail_ESS

sd(Intercept)	0.97	0.07	0.83	1.10	1.00	918	1832

Population-Level Effects:

Estimate	Est.Error	l-95% CI	u-95% CI	Rhat	Bulk_ESS	Tail_ESS

Intercept	-5.84	0.08	-5.99	-5.69	1.00	7596	8701

**as.factorStageBin2	0.71	0.05	0.61	0.81	1.00	11827	9407**

**as.factorGenoMC1R1	2.19	0.08	2.02	2.36	1.00	7040	8272**

**as.factorGenoMC1R2	2.93	0.23	2.47	3.39	1.00	7879	8678**

**as.factorGenoZcol1	0.88	0.07	0.75	1.01	1.00	8314	9244**

**as.factorGenoCorin1	0.18	0.07	0.04	0.31	1.00	7578	9057**

**as.factorGenoCorin2	0.32	0.08	0.16	0.49	1.00	6757	8399**

**as.factorGenoMC1RMUGenoZcol1	-0.31	0.11	-0.53	-0.10	1.00	7335	8280**

**as.factorGenoMC1RMUGenoZcol2	-0.79	0.30	-1.37	-0.20	1.00	8228	8810**

Family Specific Parameters:

Estimate	Est.Error	l-95% CI	u-95% CI	Rhat	Bulk_ESS	Tail_ESS

sigma	0.74	0.03	0.67	0.81	1.00	922	1892

#### Genetic architecture of spottiness

**Table S28 –** Selection of the best model explaining the spottiness in males

**Table S29** – Summary of the model best model for the genetic architecture of the spottiness in males

Family: hurdle_gamma

Links: mu = log; shape = identity; hu = logit

Formula:

SurfaceBlackSpotsCm ~ 1 + as.factor(StageBin) + as.factor(GenoZspot) + as.factor(GenoMC1R) + (1 | gr(ID, cov = grm))

hu ~ 1 + as.factor(StageBin) + as.factor(GenoZspot) + as.factor(GenoMC1R) + (1 | gr(ID, cov = grm))

Data: TabSummary[TabSummary$SexWGS == 1, ] (Number of observations: 1194)

Draws: 4 chains, each with iter = 5000; warmup = 2000; thin = 1; total post-warmup draws = 12000

Group-Level Effects: ~ID (Number of levels: 1194)

Estimate	Est.Error	l-95% CI	u-95% CI	Rhat	Bulk_ESS	Tail_ESS

sd(Intercept)	0.61	0.07	0.47	0.76	1.00	1733	3754

sd(hu_Intercept)	3.04	0.99	1.48	5.35	1.01	994	1910

Population-Level Effects:

Estimate	Est.Error	l-95% CI	u-95% CI	Rhat	Bulk_ESS	Tail_ESS

Intercept	-1.08	0.10	-1.29	-0.89	1.00	4028	6458

**hu_Intercept	-1.45	0.54	-2.66	-0.56	1.00	3075	4056**

**as.factorStageBin2	0.31	0.06	0.20	0.42	1.00	12172	8532**

**as.factorGenoZspot1	0.85	0.08	0.69	1.01	1.00	7154	8050**

**as.factorGenoZspot2	1.82	0.10	1.63	2.02	1.00	5138	7606**

**as.factorGenoMC1R1	0.57	0.07	0.44	0.70	1.00	7127	8797**

**as.factorGenoMC1R2	0.38	0.15	0.09	0.68	1.00	9962	9324**

hu_as.factorStageBin2	0.74	0.43	-0.03	1.69	1.00	5143	4451

**hu_as.factorGenoZspot1	-3.66	0.78	-5.54	-2.46	1.01	1390	2181**

**hu_as.factorGenoZspot2	-6.79	1.57	-10.46	-4.36	1.00	2258	3492**

**hu_as.factorGenoMC1R1	-3.47	0.82	-5.34	-2.13	1.01	2375	3090**

**hu_as.factorGenoMC1R2	-29.98	26.28	-101.31	-3.48	1.00	5123	4569**

Family Specific Parameters:

Estimate	Est.Error	l-95% CI	u-95% CI	Rhat	Bulk_ESS	Tail_ESS

shape	1.67	0.10	1.49	1.88	1.00	2369	5242

**Table S30 –** Selection of the best model explaining the spottiness in females

**Table S31** – Summary of the model best model for the genetic architecture of spottiness in females

Family: hurdle_gamma

Links: mu = log; shape = identity; hu = logit

Formula:

SurfaceBlackSpotsCm ~ 1 + as.factor(StageBin) + GenoZspot + GenoMC1R + (1 | gr(ID, cov = grm))

hu ~ 1 + as.factor(StageBin) + GenoZspot + GenoMC1R + (1 | gr(ID, cov = grm))

Data: TabSummary[TabSummary$SexWGS == 2, ] (Number of observations: 1286)

Draws: 4 chains, each with iter = 10000; warmup = 5000; thin = 1; total post-warmup draws = 20000

Group-Level Effects: ~ID (Number of levels: 1286)

Estimate	Est.Error	l-95% CI	u-95% CI	Rhat	Bulk_ESS	Tail_ESS

sd(Intercept)	0.47	0.05	0.37	0.56	1.00	2806	6109

sd(hu_Intercept)	1.22	0.95	0.05	3.54	1.00	2778	2353

Population-Level Effects:

Estimate	Est.Error	l-95% CI	u-95% CI	Rhat	Bulk_ESS	Tail_ESS

Intercept	0.18	0.04	0.10	0.27	1.00	9816	14007

**hu_Intercept	-4.98	0.97	-7.35	-3.59	1.00	4250	2535**

**as.factorStageBin2	0.31	0.04	0.23	0.38	1.00	25605	14287**

**GenoZspot	0.85	0.04	0.77	0.94	1.00	14683	15165**

GenoMC1R	-0.08	0.04	-0.15	0.00	1.00	20307	14499

hu_as.factorStageBin2	1.01	0.71	-0.26	2.51	1.00	15558	8570

**hu_GenoZspot	-2.59	1.03	-4.99	-0.92	1.00	17052	9081**

**hu_GenoMC1R	-0.89	0.79	-2.66	0.46	1.00	21779	11906**

Family Specific Parameters:

Estimate	Est.Error	l-95% CI	u-95% CI	Rhat	Bulk_ESS	Tail_ESS

shape	2.86	0.17	2.55	3.21	1.00	3629	7564

### Characterizing evolutionary process driving allelic and phenotypic changes

#### Observed allelic frequency changes

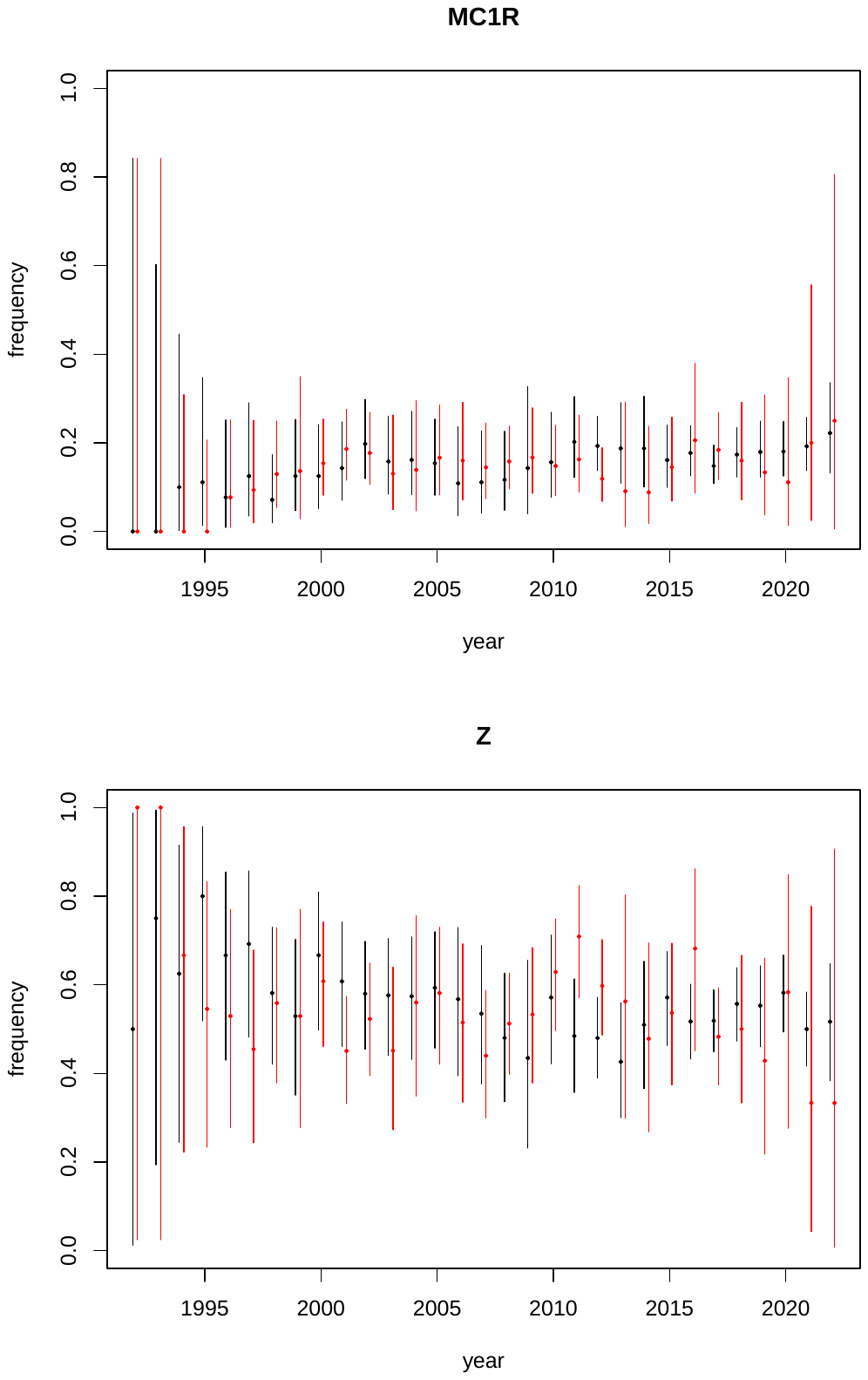

**Figure S23** – Evolution of the allelic frequencies in residents (black) and migrants (red) across the years for the major effect loci for the coloration (namely MC1R, upper panel) and Spottiness (namely Z, lower panel). Diamond depicts the estimate while vertical bars represent the confidence interval.

**Table S32** - Summary of the model of allelic frequency change as a function of year for MC1R

Call:

lm(formula = FullFreqMC1R[Years > 1995] ~ Years[Years > 1995])

Residuals:

Min	1Q	Median	3Q	Max

-0.041945 -0.012034 -0.001173 0.010482 0.051237

Coefficients:

Estimate Std. Error t value Pr(>|t|)

(Intercept)	-5.4395163 1.0171266 -5.348 1.52e-05 ***

**Years[Years > 1995] 0.0027848 0.0005063	5.500 1.03e-05 *****

---

Signif. codes: 0 ‘***’ 0.001 ‘**’ 0.01 ‘*’ 0.05 ‘.’ 0.1 ‘ ’ 1

Residual standard error: 0.02049 on 25 degrees of freedom

Multiple R-squared: 0.5475,	Adjusted R-squared: 0.5295

F-statistic: 30.25 on 1 and 25 DF, p-value: 1.027e-05

**Table S33** - Summary of the model of allelic frequency change as a function of year for Z

Call:

lm(formula = FullFreqZ[Years > 1995] ~ Years[Years > 1995])

Residuals:

Min	1Q	Median	3Q	Max

-0.079542 -0.027526 -0.005824 0.026571 0.070911

Coefficients:

Estimate Std. Error t value Pr(>|t|)

(Intercept)	4.9215914 1.9888035	2.475	0.0205 *

**Years[Years > 1995] -0.0021796 0.0009899 -2.202	0.0371 ***

---

Signif. codes: 0 ‘***’ 0.001 ‘**’ 0.01 ‘*’ 0.05 ‘.’ 0.1 ‘ ’ 1

Residual standard error: 0.04007 on 25 degrees of freedom

Multiple R-squared: 0.1624,	Adjusted R-squared: 0.1289

F-statistic: 4.848 on 1 and 25 DF, p-value: 0.03713

#### Evolutionary forces driving the frequency changes

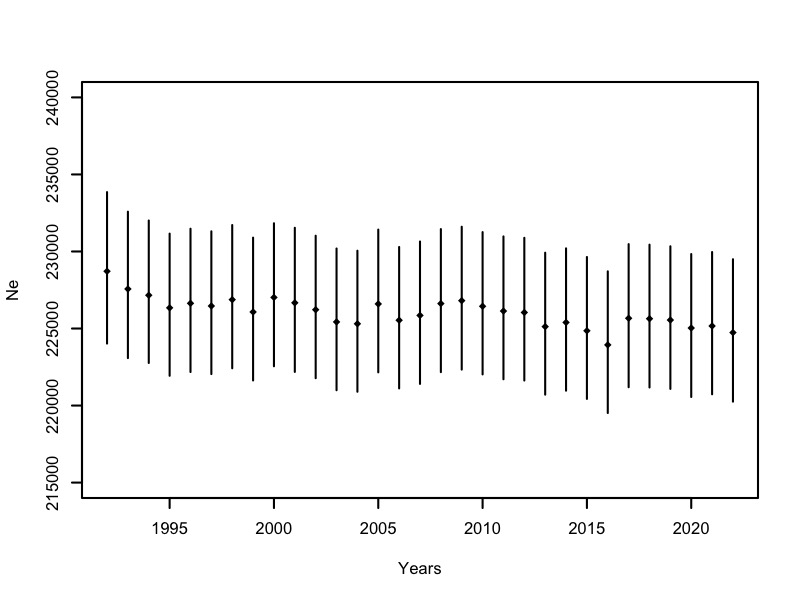

**Figure S24** – Effective population size (*Ne*) of the Swiss population across the study period. For each year, *Ne* was estimated using the adults alive that year.

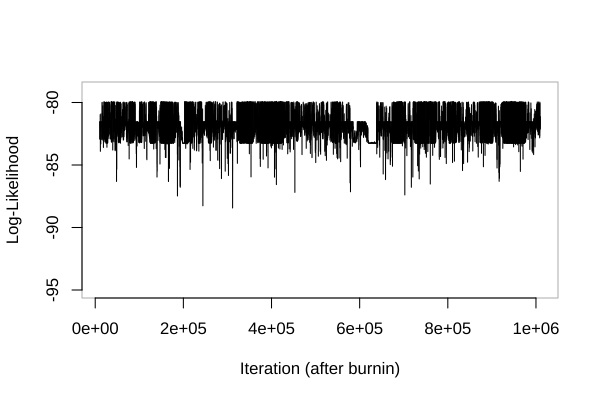

**Figure S25** – Log-likelihood trace along the MCMC chain for the model allowing jumps between selection and no-selection states at the MC1R locus, using ApproxWF.

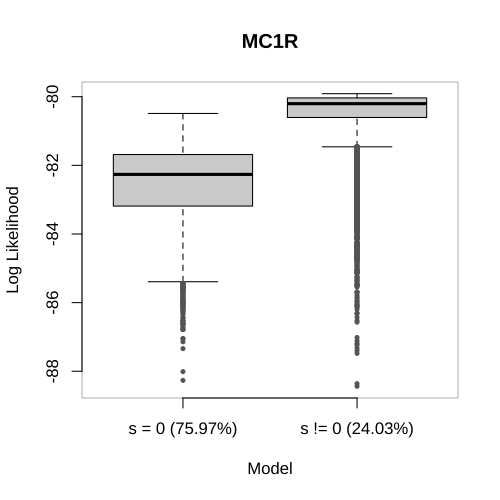

**Figure S26** – Boxplot showing the distribution of log-likelihood values for two models: selection (s!=0) and no-selection (s=0) at the MC1R locus. Values in brackets indicate the proportion of time the chain spent in each model.

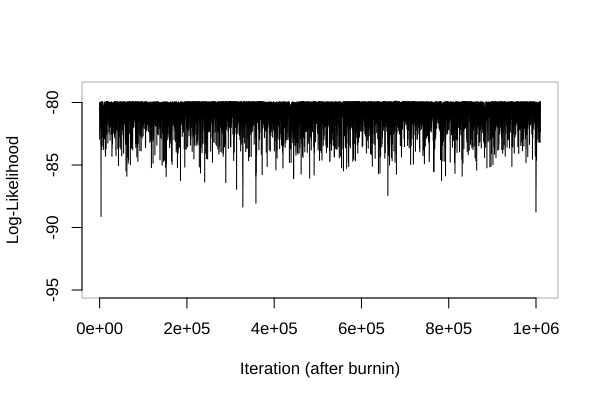

**Figure S27** – Log-likelihood trace along the MCMC chain for the model restricted to the selection state at the MC1R locus, using ApproxWF.

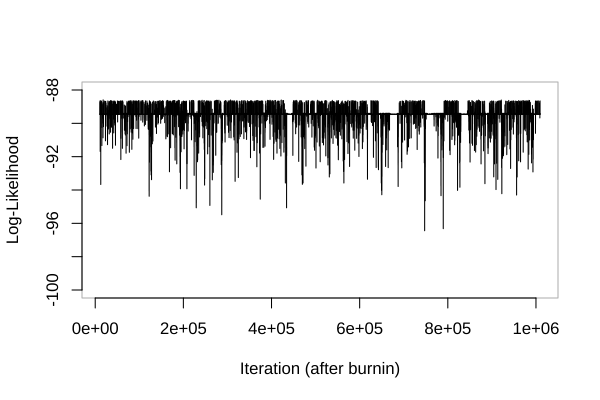

**Figure S28** – Log-likelihood trace along the MCMC chain for the model allowing jumps between selection and no-selection states at the Z locus, using ApproxWF.

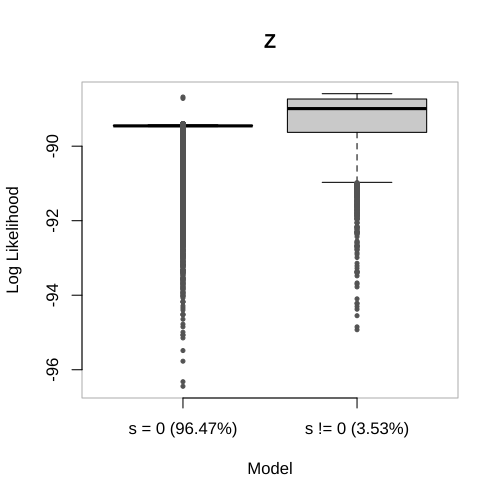

**Figure S29** – Boxplot showing the distribution of log-likelihood values for two models: selection (s!=0) and no-selection (s=0) at the Z locus. Values in brackets indicate the proportion of time the chain spent in each model.

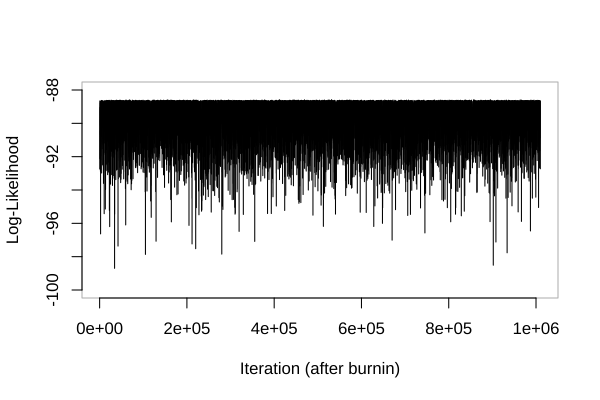

**Figure S30** – Log-likelihood trace along the MCMC chain for the model restricted to the selection state at the Z locus, using ApproxWF.

### Acknowledgments for Sample Collection

We are deeply grateful to all the individuals who generously contributed their time and effort to collecting samples over the past 30 years. This work would not have been possible without their dedication and support. The following list includes the names of those who assisted; however, it may not be entirely complete. We sincerely apologize for any unintentional omissions and extend our heartfelt thanks to everyone who involved during field work and sample collection.

alphabetically sorted list:

Albanese Margherita, Allemann Roxane, Amrein Martin, Antoniazza Silvain, Ançay Laurie, Arévalo Olympe, Audusseau Hélène, Bastardot Marc, Battesti Marine, Becciu Paolo, Bincteux Manon, Bissegger Silvan, Braaker Sonja, Bragoni Maeva, Burri Patricia, Béziers Paul, Büchi Uli, Bühler Roman, Calvani Morgane, Campon Carla, Chausson Alexandre, Chevalier Florian, Chèvre Maxime, Clément Laura, Cordes Jana, Dahuron Marine, De Gianni Lara, Deillon Clara, Demaizière Zoé, Desjardins Chloé, Dessonet Justin, Dijoux Lorine, Dolivo Vassilissa, Dreiss Amélie, Ducouret Pauline, Ducret Valérie, Dugas Mélanie, Déturche Marie, Edmé Anaïs, Ehinger Jonathan, Fioravanti Céline, Fischer Simon, Flint Raphaelle, Florent Gaëlle, Forestier Auréline, Fougère Maëlle, Frey Caroline, Freyburger Maxime, Gaillard Maryline, Gaime Florence, San-Jose Garcia Luis, Gaudy Ladina, Gelle Nathan, Gellé Nathan, Gerber Aline, Gervasio Giulia, Graf Valentin, Grangier-Bijou Anna, Gremion Jérémy, Grognuz Vincent, Guilbaut Emy, Gunter Jodok, Gémard Charlène, Heintz Anne-Caroline, Henry Isabelle, Hernandez Lucile, Hertach Matthias, Huulas Lisa, Häfliger Manuela, Ioset Noemie, Jamet Jessica, Jeanneret Yann, Judes Clarisse, Juillerat Juliette, Junker Kilian, Karsegard Charlotte, Kilchoer Noémie, Koller Andreia, Konrad Maureen, Kotur Mia, König Pascal, Küttel Stefan, Laurent Thomas, Laurent Aurélie, Le Grand Luc, Lejeune Ilou, Letuppe Thomas, Luisier Célestin, Machado Ana Paula, Marcadet Alexis, Massa Carolina, Mathis Felicia, Milliet Estelle, Morgenthaler Annick, Moulin Nicolas, Nicole Sidonie, Noll Mélissa, Nuber Maria, Oberli Frédéric, Oliveira-Xavier Aymeric, Orelli Fabien, Pedraita Serena, Pellegrini Ester, Perroud Giulia, Pilati Marco, Pini Jonathan, Pittet Jeanne, Plancherel Céline, Poncet Lisa, Questiaux Anastasia, Quetel Tiphaine, Ramseier Deborah, Rau Rebecca, Reichhart Birgit, Repnik Leona, Rieser Andreas, Robadey Aurélien, Roidt Myriam, Roland Alexandre, Romano Andrea, Rossier Virginie, Roulet Adrien, Ruppli Charlène, Rusca Arianna, Rüttimann Silvan, Sager Ramon, Sahli Christophe, Salin Kathleen, Schalcher Kim, Schmid Baptiste, Scriba Madeleine, Sironi Nicolas, Sonnay Caroline, Spori Carole, Stier Kim, Ston Daniel, Streit Jonas, Séchaud Robin, Théro Héloïse, Unger Marc, Van den Brink Valentijn, Vasseur Julien, Vermunt Aurélie, Villain Nicolas, Wassef Jérôme, Wolf Franziska, Zurkinden Damien, Zurkinden Steve.
